# Reintroduction of resistant frogs facilitates landscape-scale recovery in the presence of a lethal fungal disease

**DOI:** 10.1101/2023.05.22.541534

**Authors:** Roland A. Knapp, Mark Q. Wilber, Allison Q. Byrne, Maxwell B. Joseph, Thomas C. Smith, Andrew P. Rothstein, Robert L. Grasso, Erica Bree Rosenblum

## Abstract

Vast alteration of the biosphere by humans is causing a sixth mass extinction, driven in part by an increase in emerging infectious diseases. The emergence of the lethal fungal pathogen (*Batrachochytrium dendrobatidis*; “Bd”) has devastated global amphibian biodiversity, with hundreds of species experiencing declines or extinctions. With no broadly applicable methods available to reverse these impacts in the wild, the future of many amphibians appears grim. The once-common mountain yellow-legged (MYL) frog is emblematic of amphibians threatened by Bd. Although most MYL frog populations are extirpated following disease outbreaks, some persist and eventually recover. Frogs in these recovering populations have increased resistance against Bd infection, consistent with evolution of resistant genotypes and/or acquired immunity. We conducted a 15-year landscape-scale reintroduction study and show that frogs collected from recovering populations and reintroduced to vacant habitats can reestablish populations despite the presence of Bd. In addition, results from viability modeling suggest that many reintroduced populations have a low probability of extinction over 50 years. To better understand the role of evolution in frog resistance, we compared the genomes of MYL frogs from Bd-naive and recovering populations. We found substantial differences between these categories, including changes in immune function loci that may confer increased resistance, consistent with evolutionary changes in response to Bd exposure. These results provide a rare example of how reintroduction of resistant individuals can allow the landscape-scale recovery of disease-impacted species. This example has broad implications for the many taxa worldwide that are threatened with extinction by novel pathogens.

**Significance Statement:** Understanding how species persist despite accelerating global change is critical for the conservation of biodiversity. Emerging infectious diseases can have particularly devastating impacts, and few options exist to reverse these effects. We used large-scale reintroductions of disease-resistant individuals in an effort to recover a once-common frog species driven to near-extinction by a disease that has decimated amphibian biodiversity. Introduction of resistant frogs allowed reestablishment of viable populations in the presence of disease. In addition, resistance may be at least partially the result of natural selection at specific immune function genes, which show evidence for selection in recovering populations. The evolution of resistance and reintroduction of resistant individuals could play an important role in biodiversity conservation in our rapidly changing world.

---

**H**uman activities are increasingly impacting global biodiversity (1), with important implications for ecosystem resilience and human welfare (2). One consequence of human alteration of the biosphere is an increase in emerging infectious diseases (3, 4). Such diseases pose a severe threat to wildlife populations (5), and have caused dramatic declines and extinctions in a wide range of taxa, including echinoderms, mammals, birds, and amphibians (6–9). Amphibians are experiencing particularly devastating impacts of disease due to the recent emergence and global spread of the highly virulent amphibian chytrid fungus, *Batrachochytrium dendrobatidis* (Bd) (8, 10). By one estimate, hundreds of species have experienced Bd-caused declines, and numerous susceptible taxa are extinct in the wild (8). These impacts to global biodiversity may be unprecedented, and highlight the importance of understanding mechanisms of species persistence in the presence of emerging diseases (11).

Evidence of natural recovery in the many Bd-impacted amphibian populations is surprisingly rare (for notable exceptions, see 12–14), suggesting that disease-caused declines will be difficult to reverse. This apparent low resilience to disease effects may be due to the limited ability of many amphibians to develop Bd resistance and/or tolerance, which in turn, could also lessen the effectiveness of potential Bd mitigation strategies. Following pathogen arrival in a host population, resistance (ability to limit pathogen burden) and tolerance (ability to limit the harm caused by a particular burden) are key mechanisms to reduce disease impacts (15) and facilitate population persistence and recovery (16). Host immunity and evolution both play important roles in the development of resistance and tolerance, and utilizing these factors would seem a promising approach to developing effective strategies to mitigate disease impacts in the wild (17, 18). However, several aspects of the amphibian-Bd system present difficult obstacles, including (i) the general inability of amphibians to mount an effective immune response against Bd infection (19–21), and (ii) the apparent rarity of evolution of more resistant/tolerant genotypes (but see 22, 23). These factors suggest that reintroduction of amphibians into sites to reestablish populations extirpated by Bd will often result not in population recovery, but instead in the rapid reinfection and mortality of the introduced animals and/or their progeny (24–27). If true, the future of many amphibian species threatened by Bd appears bleak.

The mountain yellow-legged (MYL) frog, composed of the sister species *Rana muscosa* and *Rana sierrae* (28), is emblematic of the global declines of amphibians caused by Bd (8). Once the most common amphibian in the high elevation portion of California’s Sierra Nevada mountains (USA, 29), during the past century this frog has disappeared from more than 90% of its historical range (28). Due to the severity of its decline and the increasing probability of extinction, both species are now listed as “endangered” under the U.S. Endangered Species Act. In the Sierra Nevada, this decline was initiated by the introduction of non-native trout into the extensive historically-fishless region (30, 31) starting in the late 1800s. The arrival of Bd in the mid-1900s and its subsequent spread (32) caused additional large-scale population extirpations (33, 34). These Bd-caused declines are fundamentally different from the fish-caused declines because fish eradication is feasible (35) and results in the rapid recovery of frog populations (36, 37). In contrast, Bd appears to persist in habitats even in the absence of amphibian hosts (38), and therefore represents a long-term alteration of invaded ecosystems that amphibians will need to overcome to reestablish populations.

Despite the catastrophic impact of Bd on MYL frogs, wherein most Bd-naive populations are extirpated following Bd arrival (33), some populations have persisted after epizootics (during which Bd infection intensity on frogs is very high, 39) and are now recovering (Figure 1) (14). Frogs in these recovering populations show reduced susceptibility to Bd infection (14), with infection intensity (“load”) on adults consistently in the low-to-moderate range (39–41). This reduced susceptibility is evident even under controlled laboratory conditions (14), indicative of host resistance against Bd infection (and not simply an effect of factors external to individual frogs, e.g., environmental conditions). In addition to frogs from recovering populations having higher resistance to Bd infection than those from naive populations, they could also have higher tolerance, but no data are currently available to evaluate this possibility. Therefore, we focus on resistance throughout this paper. The observed resistance of MYL frogs could be the result of several non-mutually exclusive mechanisms, including natural selection for more resistant genotypes (22, 23), acquired immunity (21), and/or inherent between-population differences that pre-date Bd exposure. The possible evolution of MYL frog resistance and subsequent population recovery is consistent with that expected under “evolutionary rescue”, whereby rapid evolutionary change increases the frequency of adaptive alleles and restores positive population growth (42, 43). This intriguing possibility also suggests an opportunity to expand recovery beyond the spatial scale possible under natural recovery by utilizing resistant frogs from recovering populations in reintroductions to vacant habitats (Figure 1) (41, 44).

**Fig. 1.**
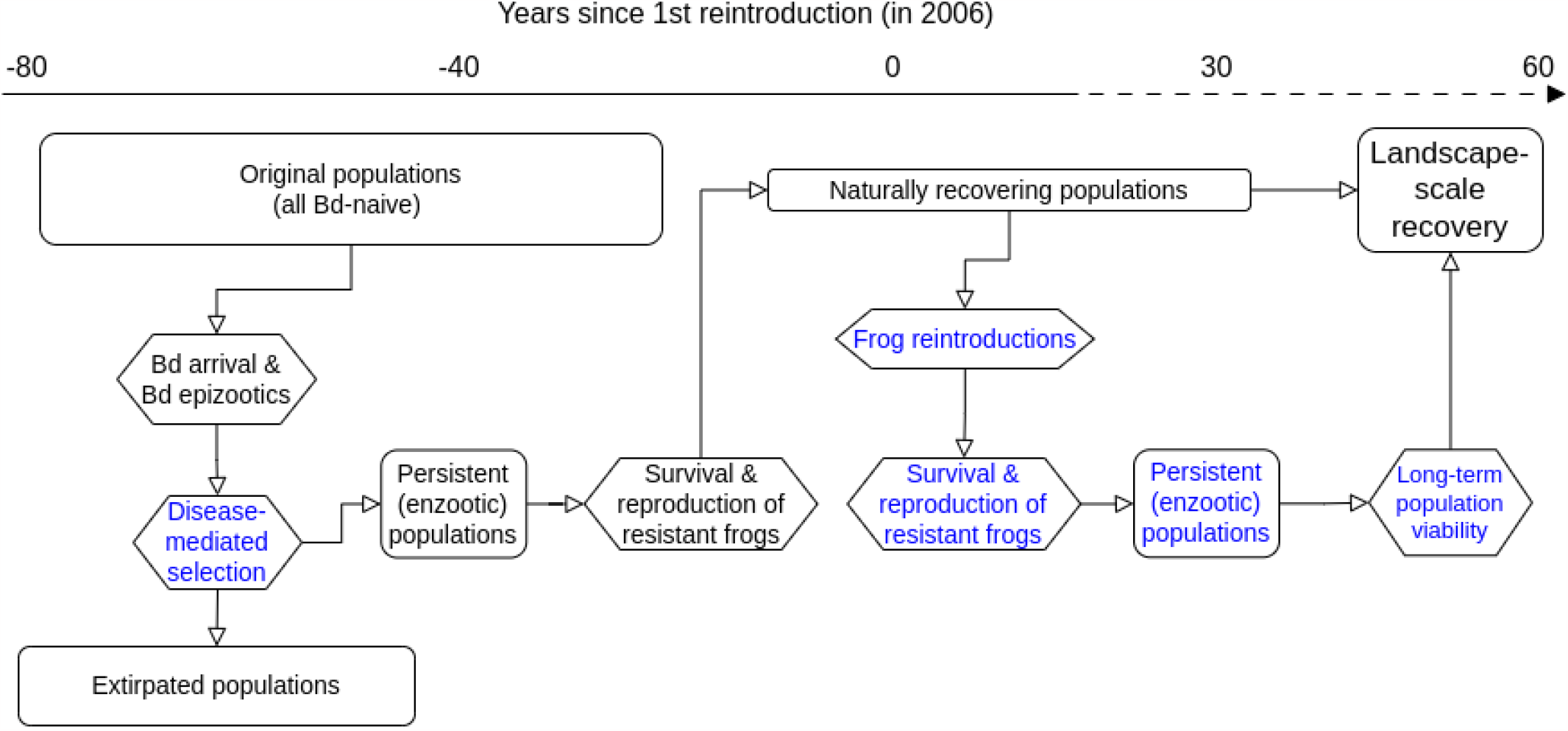
For MYL frogs, a conceptual model depicting the Bd-caused decline and subsequent natural recovery (black text), facilitated recovery via reintroductions, and the linkages between these two pathways. Rectangles and hexagons represent outcomes and processes, respectively. Blue text indicates components that are included in the current study. The timeline shows the general sequence of the components, with the dotted portion indicating a projection into the future.

In the current study, we had three primary objectives. First, to determine whether the reintroduction of resistant MYL frogs obtained from populations recovering from Bd-caused declines allows the successful reestablishment of extirpated populations despite ongoing disease, we conducted a 15-year landscape-scale frog reintroduction effort (Figure 1). Second, to extend our inferences of population recovery well beyond the temporal extent of our reintroduction study, we developed a model to estimate the probability of persistence for the reintroduced populations over a multi-decadal period (Figure 1). Third, given the importance of resistance for frog survival, population establishment, and long-term viability (this study), we conducted a genomic study using exome capture methods to determine whether MYL frogs in recovering populations show evidence of selection and whether these genomic changes are associated with resistance (Figure 1).

## Results

### Frog population recovery

To determine whether MYL frogs from recovering populations can be used to reestablish extirpated populations, we conducted 24 reintroductions in Yosemite National Park (2006–2020). Each of the reintroductions involved collection of adult frogs from 1 of 3 recovering, Bd-positive “donor” populations and translocating them to 1 of 12 nearby recipient sites (Figure S1). The donor populations included 2 of the 5 recovering populations used in the frog evolution study referenced above (Figure 5: population 1 and 4), and these donor populations contributed frogs for 20 of the 24 translocations. Following translocation, we estimated adult survival and recruitment of new adults from capture-mark-recapture (CMR) surveys and obtained counts of tadpoles and juveniles from visual encounter surveys (VES). Across all translocation sites, the duration of survey time series was 1–16 years (median = 5).

Of the 12 reintroduced populations, 9 (0.75) showed evidence of successful reproduction in subsequent years, as indicated by the presence of tadpoles and/or juveniles. For these 9 populations, one or both life stages were detected in nearly all survey-years following translocation (proportion of survey-years: median = 0.9, range = 0.29–1). These same populations were also those in which recruitment of new adults (i.e., progeny of translocated individuals) was detected. As with early life stages, recruits were detected in the majority of post-translocation survey-years (proportion of survey-years: median = 0.79, range = 0.12–1). In summary, survey results indicate that translocations resulted in the establishment of reproducing MYL frog populations at most recipient sites despite the ongoing presence of Bd.

Bd loads were fairly consistent before versus after translocation, and loads were nearly always well below the level indicative of severe chytridiomycosis (i.e., the disease caused by Bd) and associated frog mortality (Figure S2) (33, 41). Although it is possible that the observed relatively small changes in load are a consequence of individuals with high Bd loads dying and therefore being unavailable for sampling during the post-translocation period, the fact that there was little difference in pre- versus post-translocation Bd loads even in those populations that had very high frog survival (70556, 74976 - see below; Figure S2) suggests a true lack of substantial change in Bd load.

The ultimate measure of reintroduction success is the establishment of a self-sustaining population. Given that it can take years or even decades to determine the self-sustainability of a reintroduced population (for an example in MYL frogs, see 41), the use of proxies is essential for providing shorter-term insights into reintroduction success and the factors driving it. Results from our CMR surveys allowed us to accurately estimate frog survival, including over the entire CMR time series for each site and during only the 1-year period immediately following translocation. These estimates were made using site-specific models analyzed using the mrmr package. We use these estimates to describe general patterns of frog survival in all translocated cohorts, and in an among-site meta-analysis of frog survival to identify important predictors of 1-year frog survival (e.g., Bd load).

Estimates of 1-year frog survival indicate that survival was highly variable between recipient sites, but relatively constant within recipient sites (for the subset of sites that received multiple translocations; Figure 2). These patterns indicate an important effect of site characteristics on frog survival. In addition, 1-year survival was higher for frogs translocated later in the study period than earlier: 5 of the 7 populations translocated after 2013 had estimated survival *≥* 0.5, compared to only 1 of 5 populations translocated prior to 2013. We suggest this resulted primarily from our improved ability to choose recipient sites with higher habitat quality for *R. sierrae* (see **Materials and Methods - Frog population recovery - Field methods** for details). This increased survival has direct implications for population viability (see **Results - Long-term population viability**).

**Fig. 2.**
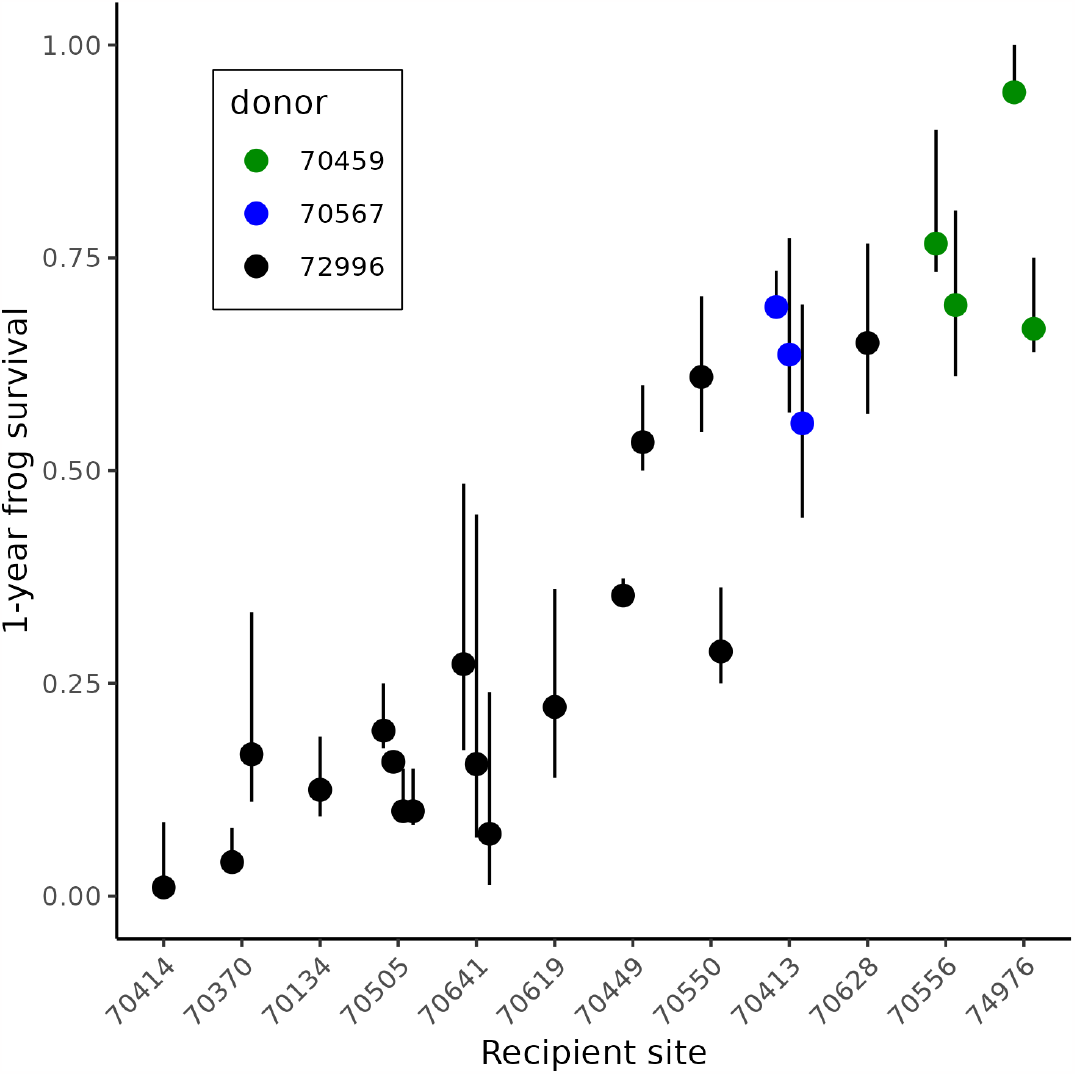
Median 1-year survival for each cohort of translocated frogs at the 12 recipient sites, as estimated for each site from the mrmr CMR model. Error bars show the 95% uncertainty intervals. Sites are arranged along the x-axis using the average of the median 1-year survival per translocation at each site. Dot colors indicate the donor population from which frogs in each translocated cohort were collected. When multiple translocations were conducted to a site, points and error bars are slightly offset to avoid overlap.

The goal of our meta-analysis was to identify important predictors of 1-year frog survival. We were particularly interested in whether Bd load had a negative effect on adult survival, as would be expected if frogs were highly susceptible to Bd infection. This analysis was conducted in a Bayesian framework and included a diversity of site, cohort, and individual-level characteristics as predictors and 1-year frog survival (Figure 2) as the response variable. The best model of 1-year frog survival identified several important predictors, but Bd load at the time of translocation was not among them (Figure S3). Instead, important predictors included winter severity in the year following translocation (snow_t1), site elevation, and donor population (Figure 3, Figure S3). Males had somewhat higher survival than females, but this effect was small (Figure 3, Figure S3). The absence of Bd load as an important predictor of frog survival is consistent with frogs in recovering populations having sufficient resistance to suppress Bd loads below harmful levels.

**Fig. 3.**
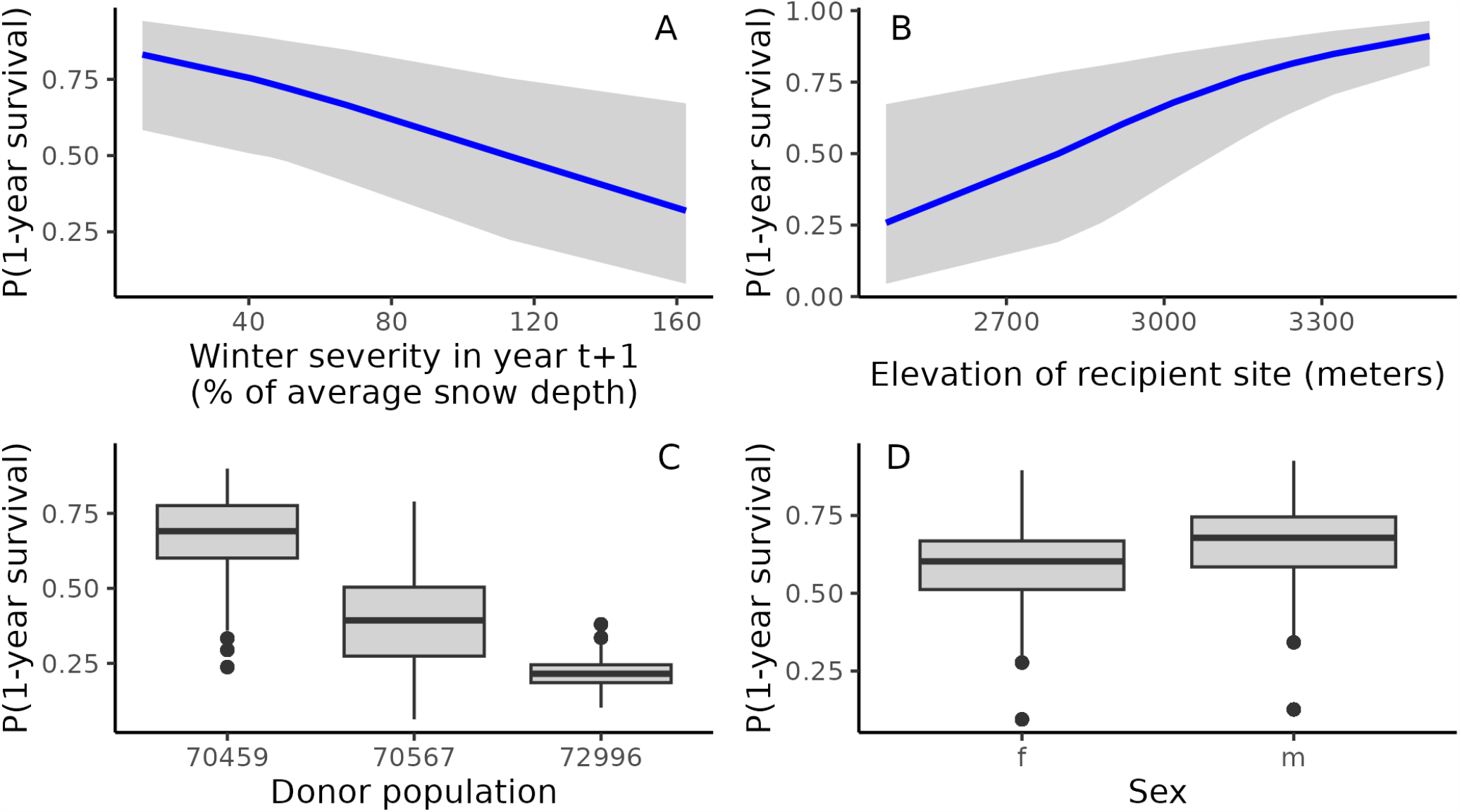
Results from the among-site meta-analysis showing conditional effects of the important predictors of 1-year frog survival (expressed as a probability): (A) winter severity in the year following translocation, (B) elevation of recipient site, (C) donor population, and (D) sex. In (A) and (B), blue lines are medians and gray ribbons are 95% uncertainty intervals. In (C) and (D), box plots show medians, first and third quartiles, largest and smallest values within 1.5x interquartile range, and values outside the 1.5x interquartile range.

In summary, results from our frog translocation study indicate that translocations resulted in (i) relatively high 1-year survival of translocated adults, as well as reproduction and recruitment, at the majority of recipient sites, (ii) 1-year survival of adults is influenced by site characteristics, weather conditions, and donor population (but not Bd load), and (iii) based on the relatively small changes in Bd load after translocation, loads appear more strongly influenced by frog characteristics (e.g., resistance) than site characteristics. Together, these results indicate that frogs translocated from recovering populations can maintain the benefits of resistance in non-natal habitats. In addition, in 3 locations where longer CMR time series allowed us to assess the survival of new adults recruited to the population, naturally-recruited adults had equivalent or higher survival probabilities than the originally translocated adults (Figure S4). This suggests that frog resistance is maintained across generations. All of the conditions described above are supportive of population establishment and long-term population growth.

### Long-term population viability

Results from the frog translocation study indicated that most populations showed evidence of successful reproduction and recruitment, and that adult survival was often relatively high (described above). Although suggestive of population establishment, a decade or more of surveys may be necessary to confirm that populations are in fact self-sustaining (41). To extend our inferences of population establishment beyond those possible from the site-specific CMR data, we developed a population viability model. Specifically, to test whether the observed yearly adult survival probabilities in translocated populations were sufficient for long-term viability, we built a stage-structured matrix model that captured known frog demography and included demographic and environmental stochasticity. We parameterized the model using CMR data from translocated populations and known life history values in this system (Table S1).

Given observed yearly adult survival probabilities of translocated frogs (from site-specific mrmr CMR models; provided in legend of Figure 4B) and a yearly survival probability of the year-1 juvenile class (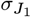) greater than 0.09, at least six of twelve translocated populations should experience a long-run growth rate *λ* greater than 1 in the presence of Bd (Figure 4A; median predicted *λ* ranges from 1.19-1.40 for these six populations). These six populations all had observed yearly adult survival greater than 0.5. As year-1 juvenile survival probability increased above 0.2, the deterministic long-run growth rate of eight of twelve population was greater than 1 (Figure 4A).

**Fig. 4.**
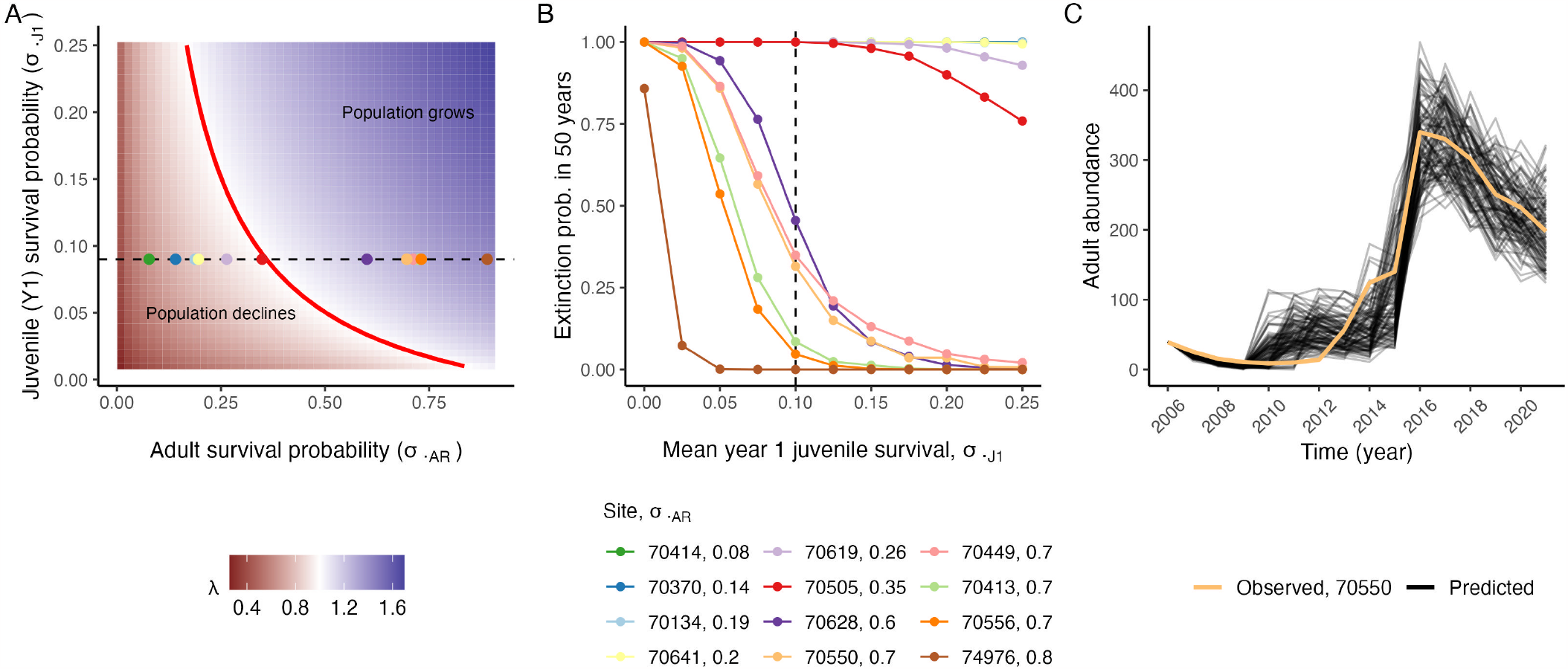
Results from population viability analysis: (A) Predicted long-run growth rate *λ* for different values of yearly adult survival probability 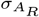 and year-1 juvenile survival and recruitment probability 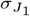, given the parameterized, deterministic model. Colored points show the predicted *λ* values for the twelve translocated populations when year-1 juvenile survival probability is 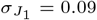 (indicated by the dashed line). The red line shows where *λ* = 1. Note that the point for 70413 is mostly hidden behind other points. (B) Predicted 50-year extinction probabilities of the 12 translocated populations, given demographic stochasticity, environmental variability in 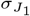, and different mean values of 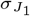. There are 6 lines at extinction probability = 1, 5 of which (70414–70619) are hidden beneath the line for 70505 when 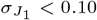. (C) 100 simulated trajectories (black lines) from the population viability model that most closely matched the observed abundance trajectory of adult amphibians at site 70550 (light orange).

**Fig. 5.**
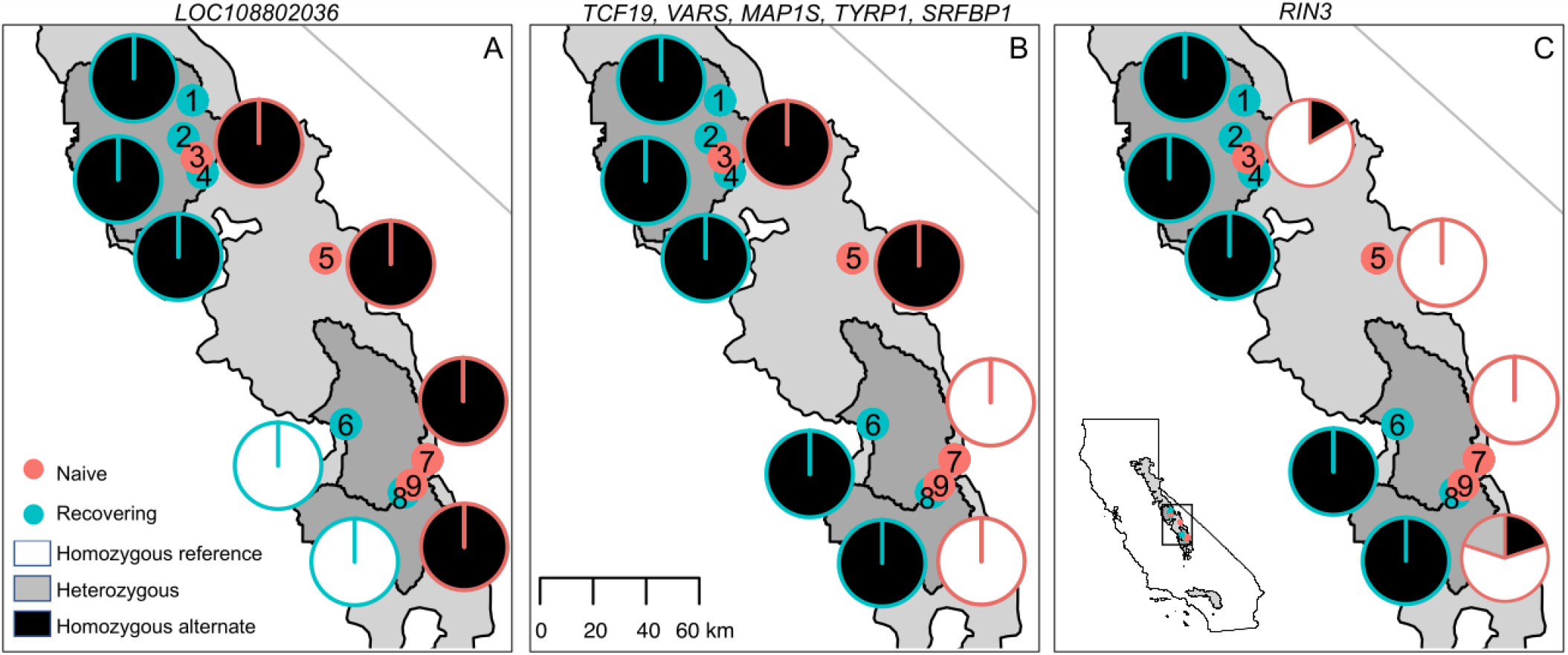
Evidence for selection on individual variants in recovering MYL frog populations at the landscape scale. For each of the 9 naive and recovering MYL frog populations (indicated by numbered points), adjacent pie charts show allele frequencies for the 11 outlier SNPs from 7 distinct genes: (A) LOC108802036, (B) TCF19, VARS, MAP1S, TYRP1, and SRFBP1, and (C) RIN3. Charts are superimposed on a map of the Sierra Nevada study area, with Yosemite, Kings Canyon, and Sequoia National Parks (from north to south) shown in dark gray, and the range boundary for MYL frogs shown in light gray. The inset map locates the study area and range boundary in California. Sites 1 and 4 (site id = 72996 and 70567, respectively) were also used as sources of frogs in the frog population recovery study.

Even when incorporating (i) demographic stochasticity and (ii) environmental stochasticity in year-1 juvenile survival and recruitment (the transition that we expect to be the most subject to environmental variability in the presence of Bd), populations with high adult survival are likely to persist over a 50 year time horizon. Our model predicted that, following a single introduction of 40 adult individuals into a population, the six populations with the highest adult survival probabilities 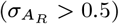 had 50-year extinction probabilities of less than 0.5 when the average year-1 juvenile survival was greater than 0.10 (Figure 4B). This indicates strong potential for long-term persistence in the presence of Bd and environmental variability in survival and recruitment. In contrast, for the six populations where yearly adult survival probability 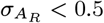, extinction probability over 50 years was always predicted to be > 50% regardless of the value of mean year-1 juvenile survival between 0 and 0.25. To test the validity of our model predictions, we demonstrated that our stochastic model could describe the general recovery trajectory of our translocated population with the longest survey history (Figure 4C; population 70550, surveyed for 16 years).

In summary, our model demonstrates that given observed yearly adult survival probabilities of translocated frogs, 50% of our translocated populations have a high probability of population growth and long-term viability in the presence of Bd. This is likely a conservative estimate because there is evidence that naturally-recruited adults have higher survival probability than translocated adults (Figure S4), but we considered these probabilities to be equal in all but three of our populations where we had sufficient data to distinguish these different probabilities.

### Frog evolution in response to Bd

Results from the preceding sections indicate the critical role of frog resistance in post-translocation frog survival, population growth, and population viability. As such, identifying the mechanisms underlying this resistance would fill a key gap in our understanding of the factors that promote population resilience in the presence of disease. Although natural selection for more resistant frog genotypes, and evolutionary rescue, may be foundational to the ability of frogs to recover despite ongoing Bd infection, for MYL frogs the role of disease-mediated selection in these processes remains unknown.

To determine whether MYL frog populations show genomic patterns consistent with an evolutionary response to Bd, we compared frog exomes (i.e., coding region of a genome) between populations with contrasting histories of Bd exposure. Specifically, we compared frog genomes sampled in 4 populations that have not yet experienced a Bd-caused epizootic (“naive”) (45) versus in 5 populations that experienced a Bd epizootic during the past several decades and have since recovered to varying degrees (“recovering”; Figure 5) (14, 33). Bd-exposure histories of the 9 study populations are based on 10-20 years of VES and Bd surveillance using skin swabbing (e.g., 14, 45, 46). Naive populations are characterized by large numbers of adults (i.e, typically 1000s), Bd prevalence that is generally 0% except during occasional Bd failed invasions (during which Bd loads remain very low, 46), and no history of Bd epizootics since we first surveyed these populations in the late 1990s and early 2000s (45). In contrast, recovering populations exist in an enzootic state (39), characterized by smaller numbers of adults (generally < 500), high Bd prevalence (often > 80%, 40), and, in adults, moderate Bd loads that are typically well below the level expected to cause mortality (33). Naive and recovering populations can be identified unambiguously using these differences in Bd prevalence and load. Finally, there is no potential for frog dispersal between the 9 study populations due to intervening distances and topography, as well as the presence of introduced (predatory) fish and fish-induced habitat fragmentation. (See **Supporting Information - Frog evolution in response to Bd - Study design** for additional details regarding the study design.)

We conducted a principal component analysis (PCA) of the genomic data to describe the relationships between sampled populations, and then used two complementary approaches to identify regions of the genome that differed between naive and recovering populations (i.e., regions under selection). First, we used a multivariate linear mixed model to evaluate associations between population type (i.e., naive versus recovering) and individual variants, including single nucleotide polymorphisms (SNPs) and insertions/deletions (INDELS), while accounting for population structure. Second, we used a splined window analysis to identify larger genomic regions showing differences between population types in *F*_*ST*_ and nucleotide diversity (*π*_*diff*_ = *π*_*naive*_ - *π*_*recovering*_).

Individual frogs clustered into 3 separate groups in principal component space (Figure S6A), and clusters reflected the species split (i.e., *R. muscosa* versus *R. sierrae*) and the strong signature of isolation-by-distance that is characteristic of MYL frogs (47–49). Importantly, each cluster contained at least one population from both the naive and recovering groups, allowing us to distinguish allelic associations of individuals sampled in the 2 population types versus allelic associations resulting from population structure and genetic drift.

Results from the individual variant and splined window analyses show that recovering populations have signatures of selection on multiple regions of the genome. The analysis of individual variants identified 11 “outlier” SNPs (i.e., showing significantly different allele frequencies between naive versus recovering populations) from 7 distinct genes across 4 contigs (Figure S6B, C). One of the 7 identified genes (LOC108802036) does not have an associated annotation. For the outlier SNPs, frequency differences between the naive and recovering populations ranged from 0.41 to 0.86. Most of these SNPs showed frequency differences in only a subset of the sampled populations (Figure 5A, B), but the SNP in the RIN3 gene showed consistent differences in frequencies across all populations (Figure 5C). This is suggestive of parallel selection at this locus across multiple populations. The other 6 outlier variants showed less consistent frequency differences across the study populations, but for these we still found a statistically significant signal of selection in 2 of the 3 genetic clusters (containing populations 5–9; Figure 5, Figure S6A). Therefore, although some outlier variant associations have a more limited geographic extent than RIN3, they still describe results that suggest parallel evolutionary changes following Bd exposure.

The splined window analysis identified 33 outlier regions for *π*_*diff*_ and 58 outlier regions for *F*_*ST*_ (Figure 6A, B). Of these, 9 regions were outliers for both metrics (“shared regions”) and 2 of these shared regions also contained one or more of the outlier SNPs described above. A total of 35 annotated genes were found in the 9 shared regions. Given this large number of genes, here we focus on those with the strongest signal of selection and/or immune-related functions. The largest *π*_*diff*_, indicative of directional selection, occurred in a 163kb region on Contig19, 12.9Mb upstream of the RIN3 outlier SNP (Figure 6C). This region contains approximately 500 SNPs and one annotated gene called “interferon-induced very large GTPase 1-like” (GVINP1). Additionally, a shared outlier region on Contig1 contained two complement factor genes (C6 and C7). Interestingly, this region had a large negative *π*_*diff*_ value, consistent with balancing selection. Finally, one shared outlier region on Contig8 contained one outlier SNP (TCF19) and was within 360kb of another outlier SNP (VARS) (Figure 6D, Figure S7). This region (854kb from the beginning of the outlier window to the VARS SNP) contained a total of 8 annotated genes. In *Xenopus*, five of these genes occur in the extended major histocompatibility complex (MHC) Class I region (FLOT1, TUBB, MDC1, CCHCR1, TCF19) and three occur in the extended MHC Class III region (HSP70, LSM2, VARS) (50). Therefore, this region under selection is part of the extended MHC Class I and III complex and shows synteny with other amphibian genomes.

**Fig. 6.**
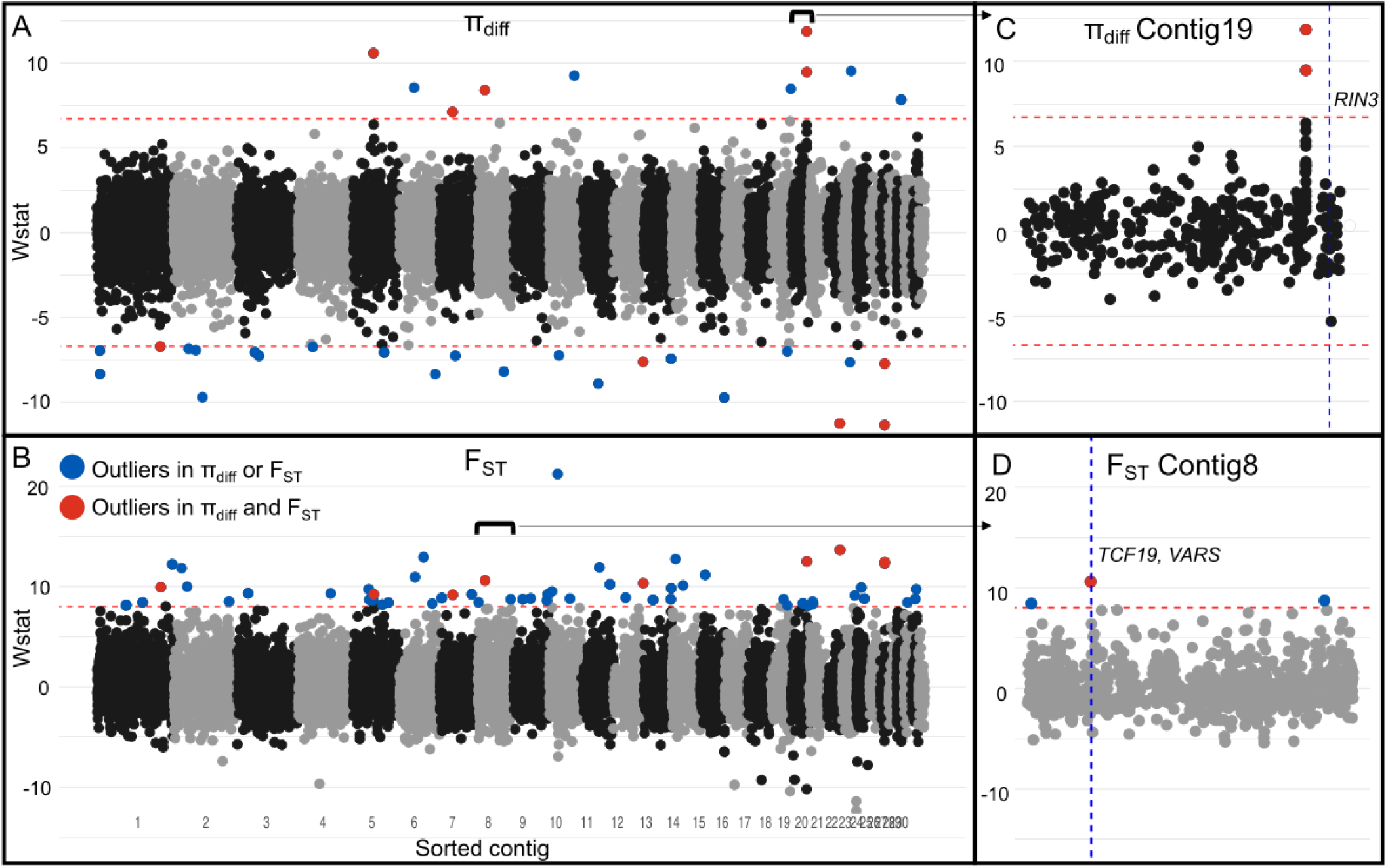
Evidence for selection on genomic regions in recovering MYL frog populations. Manhattan plot of the results from the splined window analysis showing outlier regions for the difference in (A) nucleotide diversity *π*_*diff*_ and (B) *F*_*ST*_. In (A), outlier regions are shown above the upper red dashed line and below the lower red dashed line. In (B), outlier regions are shown above the single dashed red line. Outlier regions for either *π*_*diff*_ or *F*_*ST*_ are shown in blue and outlier regions for both *π*_*diff*_ and *F*_*ST*_ are shown in red. (C) Magnified Contig19 from (A) showing two adjacent outlier regions for *π*_*diff*_ 12.9Mb upstream of the RIN3 outlier SNP (indicated with a dashed vertical blue line). (D) Magnified Contig8 from from (B) showing the *F*_*ST*_ outlier region that includes the outlier SNPs TCF19 and VARS. This region of the genome contains 8 annotated genes known to occur in the extended MHC Class I and III regions.

Although the joint processes of Bd-caused population declines and selection in response to Bd exposure could affect genetic diversity of recovering populations, we found no consistent differences in individual-level heterozygosity or population-level *π* between naive and recovering populations (see **Supporting Information - Frog evolution in response to Bd - Genetic diversity** for details). Thus, despite localized selection in particular regions of the genome, we did not find evidence for reduced genetic diversity across the genome in recovering populations. In addition, no gene ontology (GO) biological functions, molecular functions, or cellular processes were over-represented in either the outlier variants or the 35 genes located in the overlapping *F*_*ST*_ and *π*_*diff*_ splined windows (see **Supporting Information - Frog evolution in response to Bd - GO analysis** for details).

In summary, our genomic results indicate that the exomes of frogs from naive and recovering populations show substantial differences, consistent with parallel evolutionary changes following Bd exposure. The regions under selection contain several immunologically-relevant genes and gene families that are directly linked to disease resistance in other taxa.

## Discussion

Disease-induced population declines are decimating global biodiversity (5), but broadly-applicable strategies to recover affected species are generally lacking (e.g., 17). Here, we tested the possibility that populations of resistant individuals from naturally recovering populations can be used to reestablish extirpated populations of the endangered MYL frog in the presence of a highly virulent fungal pathogen (Bd). Our results indicate (i) the capacity of reintroduced populations to become established and eventually recover despite ongoing disease, (ii) that the recovering populations are likely to persist over a 50-year period, (iii) that there are substantial genomic differences between naive and recovering MYL frog populations, consistent with evolutionary change in frogs following Bd exposure, and (iv) that some of the genomic regions under selection contain genes related to disease resistance. Collectively, these results (Figure 1) provide a rare example of amphibian recovery in the presence of Bd, and have important implications for the conservation and recovery of amphibians and other taxa worldwide that are endangered by escalating impacts from emerging infectious diseases. In light of the generally low success rate of amphibian reintroduction efforts (51), our success in reestablishing MYL frog populations via translocation of resistant individuals is striking, and even more so given that MYL frogs were driven to near-extinction by Bd.

In the following discussion, we follow the sequence of frog recovery described in Figure 1 to structure our key points. Previous field studies in MYL frogs show that frog-Bd dynamics and frog survival in the presence of Bd are fundamentally different between naive and recovering populations. Following the arrival and establishment of Bd in previously-naive populations, adult frogs develop high Bd loads that lead to mass die-offs (33). In contrast, in recovering populations adult frogs typically have low-to-moderate and relatively constant Bd loads and mass die-offs are not observed (39, 40, see also Figure S2). The differences in Bd load of frogs from naive and recovering populations are also observed in controlled laboratory studies (see Figure 4 in 14), and clearly indicate that frogs from recovering populations exhibit resistance against Bd infection. This resistance could in theory be due to several factors, including natural selection for more resistant genotypes, acquired immunity, and/or inherent between-population differences that pre-date Bd exposure, but until now evidence to evaluate the role of evolution was lacking.

Results from our genomic analyses suggest that natural selection for adaptive alleles is at least partially responsible for the increased resistance of frogs in recovering populations. We identified multiple specific alleles and genomic regions showing signatures of selection between adjacent naive and recovering MYL frog populations, consistent with selection following Bd exposure. These analysis are based on samples collected from virtually all of the MYL frog populations remaining in a naive state, as well as adjacent recovering populations. This study design produced genetic clusters that each contained at least one naive and one recovering population, allowing us to detect selection without the confounding effects of population structure. In addition, we did not find a reduction in overall genetic variation in the recovering populations, suggesting that despite localized selection in the genome, these populations retain adequate genetic diversity for long-term persistence.

Importantly, some genomic regions that we identified as under selection are associated with cellular and immunological mechanisms known to contribute to disease resistance, including in amphibians (52). For example, the MHC plays an important role in immunity. In our study, we identified a region that shows evidence of selection in recovering populations and contains eight genes associated with either the MHC Class I or Class III regions. These results corroborate numerous previous studies linking MHC genes to amphibian resistance against Bd (e.g., 53, 54). Similarly, the region with the strongest indication of directional selection (as measured by *π*_*diff*_) contains the interferon-related gene GVINP1. Several previous studies of amphibians have found this gene to be differentially expressed during Bd infection (e.g., 23, 55) and in populations differing in Bd susceptibility (23). This gene is also strongly linked to disease in salmon, explaining a notable 20% of the resistance phenotype (56, 57). We also identified a region, characterized by high *F*_*ST*_ and low *π*_*diff*_, that contained the complement genes C6 and C7. The complement system plays an important role in innate immunity (58), and our results could indicate that balancing selection is acting in this region of the genome to favor a diverse set of alleles, as is known for C6 in humans (59). Based on the analysis of individual outlier variants, the RIN3 gene showed a consistent pattern of allele frequency differences across all nine of the frog populations sampled in this study, indicating consistent selection in populations distributed across a wide geographic area. This gene is associated with immune response and in *Xenopus* is expressed during appendage regeneration (60). Finally, the outlier variant with the lowest p-value was the uncharacterized gene LOC108802036. In the genome of another frog species, this gene is located adjacent to a type I interferon gene (Np-IFNi2) (61), and together with GVINP1 further suggests the importance of interferon-related genes in this system. Collectively, the genes associated with these genomic differences may confer at least some degree of resistance against Bd infection, an attribute that may be critically important to population reestablishment and recovery in the presence of Bd.

Reintroduction of resistant MYL frogs was remarkably successful in reestablishing viable populations in the presence of Bd. Of the 12 translocated populations, approximately 80% showed evidence of both successful reproduction and recruitment of new adults. Year-1 survival for 12 of the 24 translocated cohorts exceeded 50%, and > 70% of translocated cohorts had survival above this 50% level when the earliest translocations are excluded (i.e., translocations conducted when methods were still being refined; see **Materials and Methods - Frog population recovery - Field methods** for a brief description of these refinements). The fact that the relatively low Bd loads and correspondingly high frog survival was maintained when frogs were moved from donor populations to recipient sites indicates that these characteristics of naturally-recovering populations were not solely an effect of site characteristics, but were also strongly influenced by resistance inherent in the frogs. Although it could be argued that the relatively invariant Bd loads before versus after translocation are a consequence of similar pathogen pressure in the donor and translocated populations, this is at odds with the fact that in the first year after translocation frog densities are typically 1-2 orders of magnitude lower in the translocated versus donor populations and pathogen pressure should follow a similar pattern. In addition to the maintenance of Bd load and frog survival between natal and translocation sites, the relatively high survival of translocated frogs was maintained in their progeny, as expected if resistance has a genetic basis.

Results from the population viability model were also encouraging. In particular, translocated populations with > 50% survival in the first year post-translocation were predicted to have a low probability of extinction over 50 years (probability of extinction *<* 0.5 when year-1 juvenile survival probability was greater than 0.10). The viability model highlighted the important role of frog survival in affecting long-term population viability, and allowed us to extend the temporal scale of our study beyond the years covered by our post-translocation surveys. These long-term forecasts are important, given that reintroduced MYL frog populations may often take decades to achieve our ultimate goal of self-sustainability (41). Making well-supported projections about the long-term outcome of reintroduction efforts from shorter-term information is critically important to the process of adaptive management of species reintroduction programs (62), including the one we are carrying out for MYL frogs. Specifically, the combined results from our reintroduction study and viability modeling indicate that survival of frogs in the first year following translocation is an effective proxy of longer-term survival and population viability. In addition, given the repeatability of frog survival at a site, 1-year frog survival also serves as an effective proxy of site quality (i.e., the ability of a site to support high frog survival and a viable frog population over the long term). This proxy of site quality is important in the MYL frog system because accurately predicting the ability of a site to support a viable frog population *a priori* remains difficult, even after conducting 24 translocations over 16 years.

Despite the demonstrated resistance of adult MYL frogs against Bd infection, individual and population-level impacts of Bd are still evident. In an earlier study of 2 of our 12 translocated populations (41), Bd infection and load had detectable effects on the survival of adults and may have influenced population establishment (sites referred to as “Alpine” and “Subalpine” in (41) are identified as “70550” and “70505” in the current study). Applying similar analyses to all 12 of our translocated populations would likely provide a broader perspective of the ongoing effect of Bd. In addition to these important but relatively subtle effects of Bd on adults, the impacts on younger life stages are more apparent. MYL frogs immediately following metamorphosis (“metamorphs”) are highly susceptible to Bd infection (63) and as a result experience high mortality (34). This high susceptibility of metamorphs is documented in numerous species of anurans, and may result from the poorly developed immune system characteristic of this life stage (64). In naturally recovering and translocated MYL frog populations, we suggest that the high mortality of metamorphs is an important limitation on subsequent recruitment of new adults. Therefore, although adult MYL frogs appear relatively resistant, Bd infection continues to have important limiting effects on recovering populations (see also 65).

The recent emergence of Bd worldwide has contributed to the decline of hundreds of amphibian species, some of which are now extinct in the wild (8). This extraordinary impact on global amphibian biodiversity is compounded by the lack of any effective and broadly applicable strategies to reverse these impacts (17, 25). Importantly, in addition to the natural recovery documented for MYL frogs (14), other amphibian species are also showing evidence of post-epizootic recovery in the presence of Bd (12, 13) and suggest the possibility of also using animals from these recovering populations to reestablish extirpated populations. As with MYL frogs, the feasibility and long-term success of such efforts will depend on the availability of robust donor populations containing individuals that have the adaptive alleles necessary to allow frog survival and population growth in the presence of Bd. Despite the hopeful example of successful reestablishment of MYL frogs despite ongoing Bd infection, the challenge of recovering hundreds of Bd-impacted amphibian species globally is a daunting prospect. Although we now have a proven strategy to reestablish extirpated MYL frog populations, recovery across their large historical range will require substantial resources over many decades. The results of this study provide a hopeful starting point for that endeavor and other future efforts worldwide.

In our rapidly changing world, evolution is likely to play in important role in facilitating the resilience of wildlife populations. Whether the documented disease resistance in MYL frogs and concurrent recovery of decimated populations provides an airtight example of evolutionary rescue will likely always be uncertain (given that can never have a perfect understanding of the past). Regardless, we provide an example from the wild that suggests that evolution can produce individuals that harbor adaptive alleles and allow population recovery in a novel (i.e., Bd-positive) environment, and show conclusively that individuals from these recovering populations can be used to reestablish extirpated populations and expand the scale of natural recovery (Figure 1). We expect that similar species recovery actions will be an essential tool in wildlife conservation in an era of accelerating global change.

## Materials and Methods

### Frog population recovery

#### Field methods

For the 24 translocations we conducted, we identified donor populations from which adult frogs (≥ 40 mm snout-vent length) could be collected using several years of VES and skin swab collections (14), and results from population genetic analyses (48). The populations that we selected contained hundreds of *R. sierrae* adults and thousands of tadpoles. These relatively high abundances were the result of recent increases following previous Bd-caused declines (14). As is typical for recovering MYL frog populations, Bd prevalence in the donor populations was high (0.69–0.96) and Bd load (median log_10_(load) = 3.06–3.78 ITS copies) was two or more orders of magnitude below the level at which frog mortality is expected (log_10_(load) *≈* 5.78 copies)(33, 41). Recipient sites to which frogs were translocated were chosen based on previous *R. sierrae* presence (determined from VES and/or museum records) or characteristics that suggested high quality habitat for this species (66). At the beginning of this study, we had a relatively limited understanding of the factors that affect habitat quality. In subsequent years, we improved our site selection process by incorporating new information about important habitat features, in particular, overwinter habitats such as submerged boulders and overhanging banks. *R. sierrae* were absent from all recipient sites prior to the first translocation.

We conducted 1–4 translocations per site (Figure 2, Figure S1) and each translocated cohort included 18 to 99 frogs (median = 30). In preparation for each translocation, adult frogs were collected from the donor population and measured, weighed, swabbed, and PIT tagged. Frogs were transported to the recipient site either on foot or via helicopter. Following release, we visited translocated populations approximately once per month during the summer active season and conducted diurnal CMR surveys and VES (summer active season is generally July-August but can start as early as May and end as late as September; range of survey dates = May-25 to Sep-29, range of translocation dates = Jun-28 to Sep-02; median number of visits per summer = 2, range = 1–10). CMR surveys allowed estimation of adult survival, recruitment of new adults, and adult population size, and VES provided estimates of tadpole and juvenile abundance. During 2006-2012, we conducted CMR surveys on a single day (primary period) per site visit, during which we searched all habitats repeatedly for adult frogs. Frogs were captured using handheld nets, identified via their PIT tag (or tagged if they were untagged), measured, weighed, swabbed, and released at the capture location. During 2013-2022, we generally used a robust design in which all habitats were searched during several consecutive days (median number of secondary periods per primary period = 3; range = 3–7), and frogs were processed as described above. However, when the number of frogs detected on the first survey day was zero or near zero, we typically conducted only a single-day CMR survey. When using a robust design, within a primary period, frogs that were captured during more than one secondary period were measured, weighed, and swabbed during the first capture, and during subsequent captures were only identified and released.

During each site visit, we conducted VES either immediately before CMR surveys or during the first day of CMR surveys. VES was conducted by walking the entire water body perimeter, first 100 m of each inlet and outlet stream, and any fringing ponds and wetlands, and counting all *R. sierrae* tadpoles and juveniles. These *R. sierrae* life stages have high detectability, and counts are highly repeatable and provide estimates of relative abundance (31).

#### Frog counts and reproductive success

For each of the translocated populations, we used the presence of tadpoles and/or juveniles from VES and counts of new recruits (i.e, untagged adults) in CMR surveys to provide two measures of successful reproduction. To calculate the proportion of years in which tadpoles/juveniles were present at a site, we excluded surveys conducted in the year of the initial translocation to that site. This exclusion accounted for the fact that all translocations were conducted after the breeding period and reproduction would therefore not occur until the following year. Similarly, to calculate the proportion of years in which new recruits were present at a site, we excluded surveys conducted during the 3 years following the initial translocation. This accounted for the multi-year tadpole and juvenile stages in MYL frogs (Table S1).

#### Estimation of frog survival and abundance

For each translocation site, we estimated survival of translocated frogs, recruitment of new frogs into the adult population, and adult population size using a site-specific Bayesian open-population Jolly-Seber CMR model with known additions to the population (i.e., translocated cohorts), as implemented by the mrmr package (67) and using R Statistical Software (v4.4.4, 68) (see **Supporting Information - Frog population recovery - CMR model structure** for details). Briefly, the model tracks the states of *M* individuals that comprise a superpopulation made up of real and pseudo-individuals (see 41, for details). The possible states of individuals include “not recruited”, “alive”, and “dead”. The possible observations of individuals include “detected” and “not detected”. We assume that individuals that are in the “not recruited” or “dead” states are never detected (i.e., there are no mistakes in the individual PIT tag records). We also assume that new recruits were the result of within-site reproduction and not immigration from adjacent populations. This assumption is justified by the fact that no *R. sierrae* populations were present within several kilometers of the translocation sites. For all models, we used mrmr defaults for priors, number of chains (4), and warmup and post-warmup iterations (2000 for each). We evaluated convergence of the Markov chain Monte Carlo (MCMC) algorithm using trace plots and Gelman-Rubin statistics (Rhat).

#### Predictors of post-translocation frog survival

To identify important predictors of frog survival following translocation, we used multilevel Bayesian models (69, 70). Included predictor variables describe characteristics of sites, translocated cohorts, and individuals (Bd load, sex, frog size, site elevation, winter severity in the year of translocation, winter severity in the year following translocation, donor population, day of year on which a translocation was conducted, and translocation order). We used 1-year post-translocation survival estimates from CMR models as the response. Estimated survival was rounded to integer values to produce a binary outcome, and modeled with a Bernoulli distribution. Group-level (random) effects included site_id, translocation_id, or translocation_id nested within site_id. We performed all analyses with the rstanarm package (71) and R Statistical Software (v4.4.4, 68). For all models, we used default, weakly informative priors, four chains, and 5000 iterations each for warmup and post-warmup. We checked MCMC convergence using trace plots and Rhat, and evaluated model fit using leave-one-out cross-validation (72), as implemented by the loo package (73). (See **Supporting Information - Frog population recovery - Among-site survival modeling** for details.)

#### Changes in Bd load following translocation

We analyzed skin swabs using standard Bd DNA extraction and qPCR methods (74, see **Supporting Information - Frog population recovery - Laboratory methods** for details). To assess the magnitude of changes in Bd load on frogs following translocation, we compared Bd loads measured before versus after translocation. Before-translocation

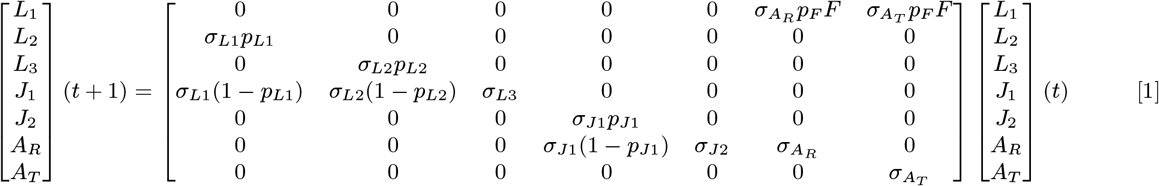

loads were quantified using skin swabs collected from all to-be-translocated frogs at the donor site on the day before or the day of the translocation. After-translocation Bd loads were based on all swabs collected from translocated frogs at the recipient site in the year of and the year following translocation. Individual frogs and their associated Bd loads were included in the dataset only if frogs were captured at the recipient site at least once during the 1-year period following translocation.

Code to replicate all of the analyses described above is available from the following GitHub repositories: https://github.com/SNARL1/translocation; https://github.com/SNARL1/cmr-analysis.

### Population viability modeling

#### Model description

To determine the implications of observed 1-year adult survival on the long-term viability of populations established via translocation, we developed a population model for MYL frogs. Our central question was: How does the magnitude and variation in observed adult survival probability across translocated populations affect the long-term persistence probability of populations? We developed a model that tracked seven state variables of a frog population: density of translocated adults (*A*_*T*_), density of adults naturally recruited into the population (*A*_*R*_), density of first-year tadpoles (*L*_1_), density of second-year tadpoles (*L*_2_), density of third-year tadpoles (*L*_3_), density of first-year juveniles (*J*_1_), and density of second-year juveniles (*J*_2_). We divided adults into two classes *A*_*T*_ and *A*_*R*_ because there is evidence that the survival probability of translocated adults and naturally recruited adults differs (Figure S4).

We modeled the dynamics of these seven state variables using a discrete-time, stage-structured model where a time step is one year. The dynamics are given by equation 1.

The parameters in this model are yearly survival probability *σ* (the subscript “” indicates a particular state variable), probability that a female frog reproduces in a given year *p*_*F*_, number of eggs produced by a female frog in a year that successfully hatch *F*, probability of a first-year tadpole remaining as a tadpole *p*_*L*1_, probability of a second-year tadpole remaining as a tadpole *p*_*L*2_, and probability of a first-year juvenile remaining as a juvenile *p*_*J*1_. First-year juvenile survival and recruitment *σ*_*J*1_ is the parameter that we think is most influenced by environmental stochasticity.

In this model we ignore density-dependent recruitment because we were interested in the growth of the population from an initial reintroduction and whether this growth was sufficient to prevent extinction over 50 years following the introduction. We also did not directly consider the dynamics of Bd in this model. We made this decision because (i) translocated populations are infected with Bd at high prevalence (41), and (ii) host density does not seem to play a significant role in multi-year Bd infection dynamics in this system (46). Thus, ignoring Bd infection dynamics and instead assuming all host vital rates are in the presence of high Bd prevalence significantly simplifies the model without much loss of realism. Additional details are provided in **Supporting Information - Population viability modeling - Incorporating yearly variability in survival rates** and **Estimating model parameters**.

#### Model analysis

After parameterizing our model with CMR-estimated adult frog survival probabilities and other known vital rates (Table S1), we performed four analyses. First, we compute the long-run growth rate *λ* for each of our 12 translocated populations to determine if the populations were deterministically predicted to grow or decline in the long-run. Second, we compute the elasticity of *λ* to four key model parameters to quantify how much changes in these parameters affected the long-run growth rate (Figure S5). This also helped us determine where in the model environmental variation in juvenile survival and recruitment would have the largest effects on population dynamics. Third, we included demographic stochasticity and environmental stochasticity in 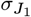 in our model and simulated the 50-year viability (i.e., 1 - extinction probability) of populations given an introduction of 40 adult individuals into an unoccupied habitat. Finally, we fit our model to our longest translocation trajectory to confirm that our model could reasonably reproduce the observed recovery trajectories of MYL frogs following reintroductions. Additional details are provided in **Supporting Information - Population viability modeling - Model analysis and simulation**. Code to replicate the analyses can be found at https://github.com/SNARL1/translocation.

### Frog evolution in response to Bd

#### Sampling and sequencing

We collected DNA samples via buccal swabbing (75) from 53 *Rana muscosa*/*Rana sierrae* individuals: 24 from 4 naive populations, and 29 from 5 recovering populations. These populations are located in the southern Sierra Nevada, from northern Yosemite National Park to northern Sequoia National Park (Figure 5). Samples were collected from 5-6 frogs per population. To minimize potential confounding effects caused by variation in frog genotypes across latitude (49), we selected sampling sites such that both population types were represented across similar latitudinal ranges. DNA was extracted following Qiagen DNEasy manufacturer’s protocols. We sequenced the samples using an exome capture assay as described in (49). Briefly, genomic libraries were prepared and captured using a custom Nimblegen capture pool. Capture baits were designed based on the coding regions of the *R. muscosa* transcriptome (GenBank accession GKCT00000000). Captured libraries were then pooled and sequenced on a NovaSeq 6000 150PE Flow Cell S1 at the Vincent J. Coates Genomics Sequencing Lab at UC Berkeley. Raw sequencing reads are available from NCBI SRA (PRJNA870451).

#### Data pre-processing and cleaning

Raw reads were filtered for adapters and contaminants using fastp (76) and aligned to the *Rana muscosa* genome (NCBI SRA: SRS6533475 (77)), with repetitive elements masked using bwa (“mem” mode) (78). Exact PCR duplicates were marked using Picard. Variants were then called following GATK best practices (v.4.2.0.0 (79)). Briefly, raw variants were called for each sample using HaplotypeCaller and combined using CombineGVCFs. Next, genotypes were jointly called using GenotypeGVCFs. Variants were then hard filtered using gatk VariantFiltration using the following parameters to remove low-quality sites: QD < 2.0, FS > 40.0, SOR > 3.0, MQ < 50.0, MQRankSum > 3.0, MQRankSum < -3.0, ReadPosRankSum > 3.0, ReadPosRankSum < -3.0. This initial filter resulted in 1,595,206 variant sites across 53 individuals.

We then further filtered our dataset at the individual and variant level. First, we trimmed our variants to only include those with minor allele frequency > 0.03, a maximum depth of 250 and minimum depth of 5, a minimum genotype quality of 20, and a maximum missing proportion of 0.5. This filter resulted in 427,038 sites, of which 353,172 were SNPs and 73,866 were INDELS. Finally, we trimmed samples with an average depth across filtered sites < 7x (n = 3). Our final dataset included 50 samples, 23 from naive and 27 from recovering populations, with an average depth of 16.7x (range = 7.4x – 26.1x).

#### Data analysis

To visualize the genomic relationships of our populations, we conducted a PCA using the glPCA function in the adegenet R package (80). To detect regions of the genome that differed between naive and recovering populations, i.e., regions under selection, we used two approaches: (1) a multivariate linear mixed model to evaluate individual variants (SNPs and INDELs), and (2) a splined window analysis to evaluate larger genomic regions. For the variant analysis, we first used a stringent data filter to include only variants with < 5% missing data (missing for no more than 2 individuals), and then calculated the likelihood ratio statistic for the resulting set of 148,307 high quality variants across 127 contigs using GEMMA (81). GEMMA calculates and incorporates a relatedness matrix for input samples, allowing us to account for relatedness and population structure when calculating likelihood ratio statistics. We identified variants showing different allele frequencies between naive versus recovering populations (“outliers”) using a Bonferroni-corrected significance level of 0.01. We visualized the results using a Manhattan plot and qqplot. We developed a more liberal set of outlier variants using a Bonferroni-corrected significance level of 0.05 and used this set solely for the gene ontology (GO) analysis (see below and **Supporting Information - Frog evolution in response to Bd - GO analysis;** Dataset S1, S2).

In the analysis of individual variants, for each outlier variant we determined whether the variant was synonymous (protein sequence the same for each variant) or non-synonymous (protein sequence differs between variants), and where in the gene it was located. To do this, we first extracted the reference genome sequence surrounding the variant using the bedtools “getfasta” function (82). Next, we re-annotated each sequence using BLAST to get the predicted gene location based on the closest annotated reference (83). We then translated each variant to amino acids and aligned this translation to that of the gene annotation to ensure proper frame of reference using Geneious (84). After ensuring proper translation, we characterized variants as within or outside the coding sequence of the gene and as either synonymous or non-synonymous.

In the splined window analysis, we identified outlier regions using *F*_*ST*_ and differences in nucleotide diversity (*π*_*diff*_) between naive and recovering populations. First, we calculated per-site *F*_*ST*_ between the naive and recovering individuals for all bi-allelic SNPs in the 30 largest contigs (98% of all SNPs) using VCFtools (85). Next, we calculated per-site nucleotide diversity *π* separately for individuals from the naive and recovering populations using VCFtools, then calculated *π*_*diff*_ for each population (*π*_*diff*_ = *π*_*naive*_ *− π*_*recovering*_). We concatenated the values for *F*_*ST*_ and *π*_*diff*_ in order of size-sorted chromosome number and adjusted the SNP position based on the relative position in the genome (for more efficient data processing and to better contextualize the strength of the outlier signals in different regions of the genome). We then used the GenWin R package (86) to conduct a splined discrete window analysis for *F*_*ST*_ and *π*_*diff*_. This method calculates where non-overlapping window boundaries should occur by identifying inflection points in the spline fitted to *F*_*ST*_ and *π*_*diff*_ values along the genome, therefore balancing false positive and false negative results that occur using other window-based methods (86). This method also calculates a W-statistic allowing for outlier identification. We identified outliers as those with a W-statistic greater than 4 standard deviations above the mean for *F*_*ST*_ or above/below the mean for *π*_*diff*_. These standard deviations represent strict criteria to select only the top ∼ 0.3% of windows. Shared outliers were then identified as those that were outliers in both analyses, meaning that they showed (i) high differentiation between naive and recovering populations, and (ii) differential patterns of nucleotide diversity in the same region. Finally, we extracted gene transcripts mapped within each region and retrieved annotation for that region using BLAST (**Supporting Information**: Datasets S3, S4).

Code used for genomic analyses and to create figures is available from the following GitHub repository: https://github.com/allie128/mylf-selection.

## ACKNOWLEDGMENTS

We thank the following for important contributions to this study: E. Hegeman and A. Lindauer–project and data management, field and laboratory work; A. Barbella and K. Rose–laboratory work; C. Kamoroff and numerous summer technicians– field work; staff at Sequoia-Kings Canyon and Yosemite National Parks, Inyo and Sierra National Forests, California Department of Fish and Wildlife, U.S. Fish and Wildlife Service, University of California-Santa Barbara Institutional Animal Care and Use Committee, and Sierra Nevada Aquatic Research Laboratory–research permits, logistical support, and/or field assistance. This project was supported by grants from the National Park Service (to R.A.K.), Yosemite Conservancy (to R.A.K.), and National Science Foundation (EF-0723563, to C. Briggs; DEB-1557190, to C. Briggs; DEB-2133401, to M.Q.W; and DBI-2120084, to C. Richards-Zawacki).

## References

1. G Ceballos, PR Ehrlich, AD Barnosky, A García, RM Pringle, TM Palmer, Accelerated modern human-induced species losses: Entering the sixth mass extinction. Sci. Adv. 1 (2015).

2. S Naeem, DE Bunker, A Hector, M Loreau, C Perrings, Biodiversity, ecosystem functioning, and human wellbeing: an ecological and economic perspective. (Oxford University Press), (2009).

3. KE Jones, NG Patel, MA Levy, A Storeygard, D Balk, JL Gittleman, P Daszak, Global trends in emerging infectious diseases. Nature 451, 990–993 (2008).

4. MC Fisher, D. Henk, CJ Briggs, JS Brownstein, LC Madoff, SL McCraw, SJ Gurr, Emerging fungal threats to animal, plant and ecosystem health. Nature 484, 186–94 (2012).

5. P Daszak, AA Cunningham, AD Hyatt, Emerging infectious diseases of wildlife – threats to biodiversity and human health. Science 287, 443–449 (2000).

6. I Hewson, JB Button, BM Gudenkauf, B Miner, AL Newton, JK Gaydos, J Wynne, CL Groves, G Hendler, M Murray,, et al., Densovirus associated with sea-star wasting disease and mass mortality. Proc. Natl. Acad. Sci. USA 111, 17278–17283 (2014).

7. MD Samuel, BL Woodworth, CT Atkinson, PJ Hart, D. LaPointe, Avian malaria in Hawaiian forest birds: infection and population impacts across species and elevations. Ecosphere 6, art104 (2015).

8. BC Scheele, F Pasmans, LF Skerratt, L Berger, A Martel, W Beukema, AA Acevedo, PA Burrowes, T Carvalho, A Catenazzi, I De la Riva, MC Fisher, SV Flechas, CN Foster, P Frías-Álvarez, TWJ Garner, B Gratwicke, JM Guayasamin, M Hirschfeld, JE Kolby, TA Kosch, E La Marca, DB Lindenmayer, KR Lips, AV Longo, R Maneyro, CA McDonald, J Mendelson, P Palacios-Rodriguez, G Parra-Olea, CL Richards-Zawacki, M. Rödel, SM Rovito, C Soto-Azat, LF Toledo, J Voyles, C Weldon, SM Whitfield, M Wilkinson, KR Zamudio, S Canessa, Amphibian fungal panzootic causes catastrophic and ongoing loss of biodiversity. Science 363, 1459–1463 (2019).

9. CX Cunningham, S Comte, H McCallum, DG Hamilton, R Hamede, A Storfer, T Hollings, M Ruiz-Aravena, DH Kerlin, BW Brook, G Hocking, ME Jones, Quantifying 25 years of disease-caused declines in Tasmanian devil populations: host density drives spatial pathogen spread. Ecol. Lett. 24, 958–969 (2021).

10. JA Luedtke, et al., Ongoing declines for the world’s amphibians in the face of emerging threats. Nature 622, 308–314 (2023).

11. RE Russell, GV DiRenzo, JA Szymanski, KE Alger, EH Grant, Principles and mechanisms of wildlife population persistence in the face of disease. Front. Ecol. Evol. 8, 569016 (2020).

12. BC Scheele, LF Skerratt, LF Grogan, DA Hunter, N Clemann, M McFadden, D Newell, CJ Hoskin, GR Gillespie, GW Heard, L Brannelly, AA Roberts, L Berger, After the epidemic: Ongoing declines, stabilizations and recoveries in amphibians afflicted by chytridiomycosis. Biol. Conserv. 206, 37–46 (2017).

13. J Voyles, DC Woodhams, V Saenz, AQ Byrne, R Perez, G Rios-Sotelo, MJ Ryan, MC Bletz, FA Sobell, S McLetchie, L Reinert, EB Rosenblum, LA Rollins-Smith, R Ibáñez, JM Ray, EJ Griffith, H Ross, CL Richards-Zawacki, Shifts in disease dynamics in a tropical amphibian assemblage are not due to pathogen attenuation. Science 359, 1517–1519 (2018).

14. RA Knapp, GM Fellers, PM Kleeman, DAW Miller, VT Vredenburg, EB Rosenblum, CJ Briggs, Large-scale recovery of an endangered amphibian despite ongoing exposure to multiple stressors. Proc. Natl. Acad. Sci. USA 113, 11889–11894 (2016).

15. L Råberg, AL Graham, AF Read, Decomposing health: tolerance and resistance to parasites in animals. Philos. Transactions Royal Soc. B: Biol. Sci. 364, 37–49 (2008).

16. LA Brannelly, HI McCallum, LF Grogan, CJ Briggs, MP Ribas, M Hollanders, T Sasso, M Familiar López, DA Newell, AM Kilpatrick, Mechanisms underlying host persistence following amphibian disease emergence determine appropriate management strategies. Ecol. Lett. 24, 130–148 (2021).

17. TWJ Garner, BR Schmidt, A Martel, F Pasmans, E Muths, AA Cunningham, C Weldon, MC Fisher, J Bosch, Mitigating amphibian chytridiomycoses in nature. Philos. Transactions Royal Soc. B: Biol. Sci. 371, 20160207 (2016).

18. DC Woodhams, J Bosch, CJ Briggs, S Cashins, LR Davis, A Lauer, E Muths, R Puschendorf, BR Schmidt, B Sheafor, J Voyles, Mitigating amphibian disease: strategies to maintain wild populations and control chytridiomycosis. Front. Zool. 8, 8 (2011).

19. EB Rosenblum, TJ Poorten, M Settles, GK Murdoch, Only skin deep: shared genetic response to the deadly chytrid fungus in susceptible frog species. Mol. Ecol. 21, 3110–3120 (2012).

20. JS Fites, JP Ramsey, WM Holden, SP Collier, DM Sutherland, LK Reinert, AS Gayek, TS Dermody, TM Aune, K Oswald-Richter, LA Rollins-Smith, The invasive chytrid fungus of amphibians paralyzes lymphocyte responses. Science 342, 366–369 (2013).

21. LF Grogan, J Robert, L Berger, LF Skerratt, BC Scheele, JG Castley, D. Newell, HI McCallum, Review of the amphibian immune response to chytridiomycosis, and future directions. Front. Immunol. 9, 2536 (2018).

22. AE Savage, KR Zamudio, Adaptive tolerance to a pathogenic fungus drives major histocompatibility complex evolution in natural amphibian populations. Proc. Royal Soc. B: Biol. Sci. 283, 20153115 (2016).

23. LF Grogan, SD Cashins, LF Skerratt, L Berger, MS McFadden, P Harlow, DA Hunter, BC Scheele, J Mulvenna, Evolution of resistance to chytridiomycosis is associated with a robust early immune response. Mol. Ecol. 27, 919–934 (2018).

24. TT Hammond, MJ Curtis, L. Jacobs, PM Gaffney, MM Clancy, RR Swaisgood, DM Shier, Overwinter behavior, movement, and survival in a recently reintroduced, endangered amphibian, Rana muscosa. J. for Nat. Conserv. 64, 126086 (2021).

25. RA Knapp, MB Joseph, TC Smith, EE Hegeman, VT Vredenburg, JE Erdman Jr, DM Boiano, AJ Jani, CJ Briggs, Effectiveness of antifungal treatments during chytridiomycosis epizootics in populations of an endangered frog. PeerJ 10, e12712 (2022).

26. M Stockwell, S Clulow, J Clulow, M Mahony, The impact of the Amphibian Chytrid Fungus on a Green and Golden Bell Frog litoria aurea reintroduction program at the Hunter Wetlands Centre Australia in the Hunter Region of NSW. Aust. Zool. 34, 379–386 (2008).

27. BC Scheele, M Hollanders, EP Hoffmann, D. Newell, DB Lindenmayer, M McFadden, DJ Gilbert, LF Grogan, Conservation translocations for amphibian species threatened by chytrid fungus: A review, conceptual framework, and recommendations. Conserv. Sci. Pract. 3, e524 (2021).

28. VT Vredenburg, R Bingham, R Knapp, JA Morgan, C Moritz, D Wake, Concordant molecular and phenotypic data delineate new taxonomy and conservation priorities for the endangered mountain yellow-legged frog. J. Zool. 271, 361–374 (2007).

29. J Grinnell, TI Storer, Animal Life in the Yosemite. (University of California Press), (1924).

30. DF Bradford, Allotopic distribution of native frogs and introduced fishes in high Sierra Nevada lakes of California: implication of the negative effect of fish introductions. Copeia 1989, 775–778 (1989).

31. RA Knapp, KR Matthews, Non-native fish introductions and the decline of the mountain yellow-legged frog from within protected areas. Conserv. Biol. 14, 428–438 (2000).

32. VT Vredenburg, SV McNally, H Sulaeman, HM Butler, T Yap, MS Koo, DS Schmeller, C Dodge, T Cheng, G Lau, CJ Briggs, Pathogen invasion history elucidates contemporary host pathogen dynamics. PLOS ONE 14, e0219981 (2019).

33. VT Vredenburg, RA Knapp, TS Tunstall, CJ Briggs, Dynamics of an emerging disease drive large-scale amphibian population extinctions. Proc. Natl. Acad. Sci. USA 107, 9689–9694 (2010).

34. LJ Rachowicz, RA Knapp, JA Morgan, MJ Stice, VT Vredenburg, JM Parker, CJ Briggs, Emerging infectious disease as a proximate cause of amphibian mass mortality. Ecology 87, 1671–1683 (2006).

35. RA Knapp, KR Matthews, Eradication of nonnative fish by gill netting from a small mountain lake in California. Restor. Ecol. 6, 207–213 (1998).

36. RA Knapp, DM Boiano, VT Vredenburg, Removal of nonnative fish results in population expansion of a declining amphibian (mountain yellow-legged frog, Rana muscosa). Biol. Conserv. 135, 11–20 (2007).

37. VT Vredenburg, Reversing introduced species effects: Experimental removal of introduced fish leads to rapid recovery of a declining frog. Proc. Natl. Acad. Sci. USA 101, 7646–7650 (2004).

38. S Walker, M Baldi Salas, D Jenkins, T Garner, A Cunningham, A Hyatt, J Bosch, M Fisher, Environmental detection of Batrachochytrium dendrobatidis in a temperate climate. Dis. Aquatic Org. 77, 105–112 (2007).

39. CJ Briggs, RA Knapp, VT Vredenburg, Enzootic and epizootic dynamics of the chytrid fungal pathogen of amphibians. Proc. Natl. Acad. Sci. USA 107, 9695–9700 (2010).

40. RA Knapp, CJ Briggs, TC Smith, JR Maurer, Nowhere to hide: impact of a temperature-sensitive amphibian pathogen along an elevation gradient in the temperate zone. Ecosphere 2, art93 (2011).

41. MB Joseph, RA Knapp, Disease and climate effects on individuals drive post-reintroduction population dynamics of an endangered amphibian. Ecosphere 9, e02499 (2018).

42. SM Carlson, CJ Cunningham, PA Westley, Evolutionary rescue in a changing world. Trends Ecol. & Evol. 29, 521–530 (2014).

43. CL Searle, MR Christie, Evolutionary rescue in host-pathogen systems. Evolution 75, 2948–2958 (2021).

44. JR Mendelson III, SM Whitfield, MJ Sredl, A recovery engine strategy for amphibian conservation in the context of disease. Biol. Conserv. 236, 188–191 (2019).

45. H Zhou, T Hanson, R Knapp, Marginal Bayesian nonparametric model for time to disease arrival of threatened amphibian populations. Biometrics 71, 1101–1110 (2015).

46. MQ Wilber, RA Knapp, TC Smith, CJ Briggs, Host density has limited effects on pathogen invasion, disease-induced declines and within-host infection dynamics across a landscape of disease. J. Animal Ecol. 91, 2451–2464 (2022).

47. AP Rothstein, RA Knapp, GS Bradburd, DM Boiano, CJ Briggs, EB Rosenblum, Stepping into the past to conserve the future: Archived skin swabs from extant and extirpated populations inform genetic management of an endangered amphibian. Mol. Ecol. 29, 2598–2611 (2020).

48. TJ Poorten, RA Knapp, EB Rosenblum, Population genetic structure of the endangered Sierra Nevada yellow-legged frog (Rana sierrae) in Yosemite National Park based on multi-locus nuclear data from swab samples. Conserv. Genet. 18, 731–744 (2017).

49. AQ Byrne, AP Rothstein, LL Smith, H Kania, RA Knapp, DM Boiano, CJ Briggs, AR Backlin, RN Fisher, EB Rosenblum, Revisiting conservation units for the endangered mountain yellow-legged frog species complex (Rana muscosa, Rana sierrae) using multiple genomic methods. Conserv. Genet. (2023).

50. Y Ohta, W Goetz, MZ Hossain, M Nonaka, MF Flajnik, Ancestral organization of the MHC revealed in the amphibian Xenopus. The J. Immunol. 176, 3674–3685 (2006).

51. C Dodd Jr, Amphibian conservation and population manipulation in Status and conservation of US amphibians, ed. M Lannoo. (Universiy of California Press), pp. 265–270 (2005).

52. KR Zamudio, CA McDonald, AM Belasen, High variability in infection mechanisms and host responses: a review of functional genomic studies of amphibian chytridiomycosis. Herpetologica 76, 189 (2020).

53. AE Savage, KR Zamudio, MHC genotypes associate with resistance to a frog-killing fungus. Proc. Natl. Acad. Sci. USA 108, 16705–16710 (2011).

54. A Bataille, SD Cashins, L Grogan, LF Skerratt, D Hunter, M McFadden, B Scheele, LA Brannelly, A Macris, PS Harlow, S Bell, L Berger, B Waldman, Susceptibility of amphibians to chytridiomycosis is associated with MHC class II conformation. Proc. Royal Soc. B: Biol. Sci. 282, 20143127 (2015).

55. AR Ellison, T Tunstall, GV DiRenzo, MC Hughey, EA Rebollar, LK Belden, RN Harris, R Ibáñez, KR Lips, KR Zamudio, More than skin deep: functional genomic basis for resistance to amphibian chytridiomycosis. Genome Biol. Evol. 7, 286–298 (2014).

56. D Robledo, A Hamilton, A. Gutiérrez, JE Bron, RD Houston, Characterising the mechanisms underlying genetic resistance to amoebic gill disease in Atlantic salmon using RNA sequencing. BMC Genomics 21 (2020).

57. D Robledo, O Matika, A Hamilton, RD Houston, Genome-wide association and genomic selection for resistance to amoebic gill disease in Atlantic salmon. G3 Genes|Genomes|Genetics 8, 1195–1203 (2018).

58. M Riera Romo, D Pérez-Martínez, C Castillo Ferrer, Innate immunity in vertebrates: an overview. Immunology 148, 125–139 (2016).

59. M Soejima, H Tachida, M Tsuneoka, O Takenaka, H Kimura, Y Koda, Nucleotide sequence analyses of human complement 6 (C6) gene suggest balancing selection. Annals Hum. Genet. 69, 239–252 (2005).

60. T Fukazawa, Y Naora, T Kunieda, T Kubo, Suppression of the immune response potentiates tadpole tail regeneration during the refractory period. Development 136, 2323–2327 (2009).

61. Z Gan, YC Yang, SN Chen, J Hou, ZA Laghari, B Huang, N Li, P Nie, Unique composition of intronless and intron-containing type I IFNs in the Tibetan frog Nanorana parkeri provides new evidence to support independent retroposition hypothesis for type I IFN genes in amphibians. The J. Immunol. 201, 3329–3342 (2018).

62. PJ Seddon, DP Armstrong, RF Maloney, Developing the science of reintroduction biology. Conserv. Biol. 21, 303–312 (2007).

63. S Ellison, RA Knapp, W Sparagon, A Swei, VT Vredenburg, Reduced skin bacterial diversity correlates with increased pathogen infection intensity in an endangered amphibian host. Mol. Ecol. 28, 127–140 (2019).

64. JE Humphries, C. Lanctôt, J Robert, HI McCallum, D. Newell, LF Grogan, Do immune system changes at metamorphosis predict vulnerability to chytridiomycosis? an update. Developmental and Comparative Immunology 136, 104510 (2022).

65. M Hollanders, LF Grogan, CJ Nock, HI McCallum, D. Newell, Recovered frog populations coexist with endemic Batrachochytrium dendrobatidis despite load-dependent mortality. Ecol. Appl. 33 (2022).

66. RA Knapp, Effects of nonnative fish and habitat characteristics on lentic herpetofauna in Yosemite National Park, USA. Biol. Conserv. 121, 265–279 (2005).

67. MB Joseph, mrmr: mark recapture miscellany in R (2019) R package version 0.1.1.

68. R Core Team, R: A Language and Environment for Statistical Computing (R Foundation for Statistical Computing, Vienna, Austria), (2022).

69. A Gelman, JB Carlin, HS Stern, DB Dunson, A Vehtari, DB Rubin, Bayesian Data Analysis. (CRC Press), (2013).

70. J Gabry, D Simpson, A Vehtari, M Betancourt, A Gelman, Visualization in Bayesian workflow. J. Royal Stat. Soc. Ser. A 182, 389–402 (2019).

71. B Goodrich, J Gabry, I Ali, S Brilleman, rstanarm: Bayesian applied regression modeling via Stan. (2022) R package version 2.21.3.

72. A Vehtari, A Gelman, J Gabry, Practical Bayesian model evaluation using leave-one-out cross-validation and WAIC. Stat. Comput. 27, 1413–1432 (2016).

73. A Vehtari, J Gabry, M Magnusson, Y Yao, PC Bürkner, T Paananen, A Gelman, loo: Efficient leave-one-out cross-validation and waic for Bayesian models (2022) R package version 2.5.1.

74. D Boyle, D Boyle, V Olsen, J Morgan, A Hyatt, Rapid quantitative detection of chytridiomycosis (Batrachochytrium dendrobatidis) in amphibian samples using real-time Taqman PCR assay. Dis. Aquatic Org. 60, 141–148 (2004).

75. T Broquet, L Berset-Braendli, G Emaresi, L Fumagalli, Buccal swabs allow efficient and reliable microsatellite genotyping in amphibians. Conserv. Genet. 8, 509–511 (2007).

76. S Chen, Y Zhou, Y Chen, J Gu, fastp: an ultra-fast all-in-one fastq preprocessor. Bioinformatics 34, i884–i890 (2018).

77. T Hon, K Mars, G Young, YC Tsai, JW Karalius, JM Landolin, N Maurer, D Kudrna, MA Hardigan, CC Steiner, SJ Knapp, D Ware, B Shapiro, P Peluso, DR Rank, Highly accurate long-read hifi sequencing data for five complex genomes. Sci. Data 7 (2020).

78. H Li, Aligning sequence reads, clone sequences and assembly contigs with bwa-mem (2013).

79. GA Van der Auwera, BD O’Connor, Genomics in the cloud: using Docker, GATK, and WDL in Terra. (O’Reilly Media), p. 496 (2020).

80. T Jombart, adegenet : a R package for the multivariate analysis of genetic markers. Bioinformatics 24, 1403–1405 (2008).

81. X Zhou, M Stephens, Efficient multivariate linear mixed model algorithms for genome-wide association studies. Nat. Methods 11, 407–409 (2014).

82. AR Quinlan, IM Hall, BEDTools: a flexible suite of utilities for comparing genomic features. Bioinformatics 26, 841–842 (2010).

83. S Altschul, Gapped BLAST and PSI-BLAST: a new generation of protein database search programs. Nucleic Acids Res. 25, 3389–3402 (1997).

84. M Kearse, R Moir, A Wilson, S Stones-Havas, M Cheung, S Sturrock, S Buxton, A Cooper, S Markowitz, C Duran, T Thierer, B Ashton, P Meintjes, A Drummond, Geneious Basic: An integrated and extendable desktop software platform for the organization and analysis of sequence data. Bioinformatics 28, 1647–1649 (2012).

85. P Danecek, A Auton, G Abecasis, CA Albers, E Banks, MA DePristo, RE Handsaker, G Lunter, GT Marth, ST Sherry, G McVean, R Durbin, The variant call format and VCFtools. Bioinformatics 27, 2156–2158 (2011).

86. TM Beissinger, GJ Rosa, SM Kaeppler, D Gianola, N de Leon, Defining window-boundaries for genomic analyses using smoothing spline techniques. Genet. Sel. Evol. 47, 1–9 (2015).

