## Supporting Information for "Reintroduction of resistant frogs facilitates landscape-scale recovery in the presence of a lethal fungal disease"

**This PDF file includes:**

Supporting text
Figs. S1 to S10
Table S1
Legends for Dataset S1 to S4
SI References

**Other supporting materials for this manuscript include the following:**

Datasets S1 to S4

### Supporting Information Text

#### 1. Frog population recovery

**A. Laboratory methods.** Swab extracts were analyzed using standard Bd DNA extraction and qPCR methods (1), and extracts were analyzed singly instead of in triplicate (2). For analysis of swabs collected during 2005–2014, we used standards developed from known concentrations of zoospores (1), and after 2014, we used standards based on single ITS1 PCR amplicons (3). Based on paired comparisons between samples analyzed using both types of standards, Bd in the study area has an average of 60 ITS1 copies per zoospore. To express all qPCR results as the number of ITS1 copies, starting quantities obtained using the zoospore standard (measured as "zoospore equivalents") were multiplied by 60. In addition, all qPCR quantities (regardless of standard) were multiplied by 80 to account for the fact that DNA extracts from swabs were diluted 80-fold during extraction and PCR (4).

**B. CMR model structure.** We estimated survival and recruitment for each site using open population CMR models based on (5). For each individual $i = 1, ..., M$ on each survey $j = 1, ..., n_j$: $o_{i,j} = 1$ if the individual was not detected, and $o_{i,j} = 2$ if the individual was detected. Capture histories of $M$ individuals are modeled, although only $N_s$ individuals were captured. This parameter expanded data augmentation allows us to capture the possibility that undetected individuals may have recruited into the adult population (6). Here, $M$ was chosen to be three times the number of observed individuals ($3N_s$) to be considerably greater than our prior guess of $N_s$.

We denote the true state of individual $i$ as $u_{i,t}$ for primary period $t = 1, ..., n_t$. The four states that we consider are: $u_{i,t} = 1$ for individuals that have not recruited, $u_{i,t} = 2$ for live adults, and $u_{i,t} = 3$ for dead adults. Each survey $j = 1, ..., n_j$ occurs in one of the $n_t$ primary periods, and we denote the primary period in which survey $j$ takes place as $t_j$. Each primary period $t$ occurs within one year, but within a year there can be multiple primary periods. We set the year containing the first primary period to $y_{t=1} = 1$, and generally $y_t$ represents the year containing primary period $t$. Years increment by one until the final year of the mark recapture efforts, which we denote $n_y$: $y \in \{1, 2, ..., n_y\}$. We assume that within a primary period, the state of each individual does not change (i.e., individuals do not recruit into the adult population, gain or lose Bd infection, or die). This assumption is justified by the short time intervals between surveys within primary periods, in cases where primary periods contain multiple secondary periods.

Live individuals are detected with probability $p_j$, which is modeled as:

$$p_j = \text{logit}^{-1}(X_j^{(p)} \beta^{(p)}),$$

where $X_j^{(p)}$ is a known row vector and $\beta^{(p)}$ an unknown parameter vector. Not recruited and dead individuals are never captured. We bundle these assumptions about the observation probabilities for survey $j$ into an emission matrix $\Omega_j$:

$$\Omega_j = \begin{array}{c} \\ \end{array} \begin{matrix} \text{Not detected} & \text{Detected} \\ \left(\begin{matrix} 1 \\ 1 - p_j \\ 1 \end{matrix}\right. & \left.\begin{matrix} 0 \\ p_j \\ 0 \end{matrix}\right) \end{matrix} \begin{matrix} \text{Not recruited} \\ \text{Alive} \\ \text{Dead} \end{matrix}$$

The state transition matrix $\Psi_{t,i}$ contains the probabilities of individual $i$ transitioning from state $u_{i,t}$ (rows) to $u_{i,t+1}$ (columns) between primary period $t$ and $t + 1$. For non-introduced (i.e., naturally recruited) individuals, this matrix is given by:

$$\Psi_{t,i} = \begin{matrix} \text{Not recruited} & \text{Alive} & \text{Dead} \\ \left(\begin{matrix} 1 - \lambda_t \\ 0 \\ 0 \end{matrix}\right. & \begin{matrix} \lambda_t \\ \phi_t \\ 1 \end{matrix} & \left.\begin{matrix} 0 \\ 1 - \phi_t \\ 1 \end{matrix}\right) \end{matrix} \begin{matrix} \text{Not recruited} \\ \text{Alive} \\ \text{Dead} \end{matrix},$$

where $\lambda_t$ is the probability of recruiting in time $t$ and $\phi_t$ is the probability of survival in time $t$.

For introduced individuals, which have deterministic recruitment (i.e., they recruit when introduced), the state transition matrix is given by:

$$\Psi_{t,i} = \begin{matrix} \text{Not recruited} & \text{Alive} & \text{Dead} \\ \left(\begin{matrix} 1 - I_{t,i} \\ 0 \\ 0 \end{matrix}\right. & \begin{matrix} I_{t,i} \\ \phi_t \\ 1 \end{matrix} & \left.\begin{matrix} 0 \\ 1 - \phi_t \\ 1 \end{matrix}\right) \end{matrix} \begin{matrix} \text{Not recruited} \\ \text{Alive} \\ \text{Dead} \end{matrix},$$

where $I_{t,i}$ is a known indicator function for whether individual $i$ was introduced in primary period $t$.

We allow recruitment probabilities to vary in time via random effects, such that:

$$\lambda_t = \text{logit}^{-1}(\alpha^{(\lambda)} + \epsilon_t^{(\lambda)}),$$

where $\alpha^{(\lambda)}$ is an intercept parameter and $\epsilon_t^{(\lambda)}$ is an adjustment for time $t$.

Survival probabilities also vary in time, and as a function of known covariates:

$$\phi_t = \text{logit}^{-1}(X_t^{(\phi)}\beta^{(\phi)} + \epsilon_t^{(\phi)}),$$

where $X_t^{(\phi)}$ is a row vector of known covariates, $\beta^{(\phi)}$ is a column vector of unknown coefficients, and $\epsilon_t^{(\phi)}$ is an adjustment for time $t$.

To complete the specification of the Bayesian model, we specify priors for all unknown parameters. The recruitment parameter priors were specified as follows:

$$\alpha^{(\lambda)} \sim N(0,1),$$

$$\sigma^{(\lambda)} \sim N_+(0,1),$$

$$\epsilon_t^{(\lambda)} \sim N(0,\sigma^{(\lambda)}),$$

for periods $t = 1, ..., T$. Here $N$ represents the normal distribution and $N_+$ the half normal distribution with positive support.

Survival parameter priors were specified similarly as:

$$\beta^{(\phi)} \sim N(0,1),$$

$$\sigma^{(\phi)} \sim N_+(0,1),$$

$$\epsilon_t^{(\phi)} \sim N(0,\sigma^{(\phi)}),$$

for time $t = 1, ..., T$.

The detection model coefficient vector also received a standard normal prior $\beta^{(p)} \sim N(0,1)$.

We computed the likelihood of each individual capture history using the forward algorithm, and we estimated the latent states using the forward-backward algorithm (5, 7).

All of the code to specify and fit the model in Stan is available in the open source mrmr package (8).

The joint distribution of the resulting model can be written as follows:

$$[\alpha^{(\lambda)}, \sigma^{(\lambda)}, \epsilon_{1:T}^{(\lambda)}, \beta^{(\phi)}, \sigma^{(\phi)}, \epsilon_{1:T}^{(\phi)}, \beta^{(p)} \mid \boldsymbol{Y}] \propto \prod_{i=1}^{M} [Y_i \mid \alpha^{(\lambda)}, \epsilon_{1:T}^{(\lambda)}, \beta^{(\phi)}, \epsilon_{1:T}^{(\phi)}, \beta^{(p)}] \times$$

$$\prod_{t=1}^{T} [\epsilon_t^{(\lambda)} \mid \sigma^{(\lambda)}][\epsilon_t^{(\phi])} \mid \sigma^{(\phi)}][\sigma^{(\lambda)}][\sigma^{(\phi)}][\alpha^{(\lambda)}][\beta^{(\phi)}][\beta^{(p)}],$$

where $\boldsymbol{Y}$ is an $M \times T$ detection matrix, and $Y_i$ the capture history of individual $i$.

**C. Among-site survival modeling.** The objective of this analysis is to describe the influence of site, cohort, and individual level characteristics on post-translocation frog survival. By modeling survival estimates obtained from site-specific mrmr CMR analyses, we are in effect conducting an among-site meta-analysis. Although it would theoretically be possible to estimate survival covariate effects in a joint CMR model that integrates capture histories across all sites, this was impractical due to computational requirements of the CMR models (namely, run time and memory).

We used Bayesian generalized linear mixed models to investigate predictors of survival among sites. The response $y_i$ is binary, representing a point estimate of whether individual $i$ survived in the year following translocation. We generated these point estimates by rounding the posterior median of 1-year post-introduction survival for each individual (from site-specific mrmr CMR models) and modeled the data using a Bernoulli distribution:

$$y_i \sim \text{Bernoulli}(p_i),$$

where $p_i$ is the probability of survival.

We modeled variation in probabilities as follows:

$$\text{logit}(p_i) = \alpha + X_i\boldsymbol{\beta} + \boldsymbol{\nu}_{g[1:N]},$$

where $\alpha$ is an intercept, $X_i$ is a length $K$ row vector of predictors, $\boldsymbol{\beta}$ is a column vector of predictor effects, and $\boldsymbol{\nu}$ a vector of group level random effects. Here $g[i]$ refers to the group $g$ containing individual $i$, and we estimate an adjustment for each of the $G$ groups $(\nu_1, ..., \nu_G)$.

These models were fit using the `stan_glmer()` function in the rstanarm package, with default priors described below (9). These priors are vague, but include data-dependent scaling as follows to account for different input variable scales. However, because we standardized all predictor variables similarly to have equal variance (by centering and dividing by twice the sample standard deviation), the resulting priors are identical. Specifically, we have:

$$\alpha \sim \text{Normal}(0, 2.5),$$

and

$$\beta_k \sim \text{Normal}(0, 5),$$

for $k = 1, ..., K$ where $K$ is the number of predictor variables.

The default prior for group level adjustments $\nu_1, ..., \nu_G$ in rstanarm is a zero-mean Gaussian, where the covariance matrix is constructed from a correlation matrix with an LKJ prior, and a vector of variance parameters – the decomposition of variance prior with unit regularization, concentration, shape, and scale parameters (10).

We drew posterior samples using Dynamic Hamiltonian Monte Carlo in Stan, with four parallel chains, each run for 10,000 iterations, discarding the first half of each chain as warm-up draws (9). We used Rhat statistics and trace plots to verify convergence. We considered models with different subsets of fixed and random effects, and used approximate leave-one-out cross validation to identify the best model (11).

### 2. Population viability modeling

**A. Incorporating yearly variability in vital rates.** We computed yearly survival probabilities for translocated adults $\sigma_{A_T}$ and naturally recruited adults $\sigma_{A_R}$ from the posterior distribution of individual state trajectories derived from mrmr CMR models. Although we observed yearly variability in adult survival within a population, the magnitude of this variability was small compared to among-population variability (Fig. 2). Thus, we did not include yearly within-population variability in adult survival in this analysis. However, within a population there was substantial yearly variability in the successful recruitment of adults, greater than what we would expect from Poisson variability around a mean value. Therefore, we allowed for yearly variability in the probability of year-1 juvenile survival and recruitment (additional details provided in **Estimating model parameters** below). We also could have included environmental stochasticity in year-2 juvenile survival and recruitment $\sigma_{J_2}$, but our elasticity analysis (**Supporting Information - Population viability modeling - Model analysis and simulation**) showed that this parameter had little effect on host growth rate relative to $\sigma_{J_1}$ (Figure S5).

**B. Estimating model parameters.** The baseline parameter values for the model and how they were estimated are given in Table S1. Parameters $\sigma_{A_T}$ and $\sigma_{A_R}$ were extracted directly from our CMR models (see **Materials and Methods - Frog populaton recovery - Estimation of frog survival and abundance** and **Supporting Information - Frog population recovery - CMR model structure** for details). For populations where we had a sufficient number of PIT-tagged, naturally-recruited adults, we observed that $\sigma_{A_T}$ and $\sigma_{A_R}$ could be notably different, with $\sigma_{A_R} > \sigma_{A_T}$ (Figure S4). For populations lacking sufficient numbers of naturally-recruited adults, we were unable to directly estimate $\sigma_{A_R}$, and instead set $\sigma_{A_R} = \sigma_{A_T}$.

To estimate the $\sigma_{A_R}$, we used the posterior distribution of predicted true states for naturally-recruited individuals (1=not recruited, 2=alive, 3=dead, as described in **Supporting Information - Frog population recovery - CMR model structure**), then calculated the posterior probability of individuals surviving between consecutive primary periods, conditional on being alive in the first primary period (e.g., given a value of 2 (alive) in the first primary period, how often was the value still 2 (alive) in the next primary period compared to 3 (dead)?) . This yielded posterior distributions for survival probabilities between primary periods. However, because the time interval between primary periods differed, the survival probabilities between different consecutive primary periods were not directly comparable. To address this, we converted the survival probabilities between each consecutive primary period to per day death rates, propagating the uncertainty from the posterior distributions. We then took a weighted average of these death rates, weighted by the time interval between primary periods, to get the average per day death rate over the entire CMR survey. We converted this per day death rate $d$ back to a yearly survival probability using $\exp(-d \times 365 \text{ days})$. We used the same procedure for $\sigma_{A_T}$ such that our estimates of average yearly survival probability were comparable between $\sigma_{A_R}$ and $\sigma_{A_T}$.

**C. Model analysis and simulation.** We performed four analyses on our model. First, we considered a deterministic version of our model with no yearly heterogeneity in year-1 juvenile survival and recruitment probability $\sigma_{J_1}$, and calculated the predicted long-run growth rate $\lambda$ of a population for different values of $\sigma_{A_R}$ and $\sigma_{J_1}$. We then fixed $\sigma_{J_1} = 0.09$ and calculated the predicted growth rate of our 12 populations.

Second, we performed a local elasticity analysis on $\lambda$ with respect to parameters $\sigma_{J_1}$, $\sigma_{J_2}$, $\sigma_{A_R}$, and $F$ to determine how small changes in these parameters could influence the long-run deterministic growth rate of populations (Figure S5).

Third, we defined a version of the model with demographic and environmental stochasticity, where environmental stochasticity was represented by among-year variability in $\sigma_{J_1}$. We used this model to simulate a one-time introduction of 40 translocated adult frogs. We ran this simulation 1000 times for each population and computed the probability of a population becoming extinct after 50 years given the observed parameter values and environmental stochasticity in $\sigma_{J_1}$. We varied the mean recruitment probability $\sigma_{J_1}$ from 0 and 0.25 and drew values of $\sigma_{J_1}$ each year from a beta distribution with a dispersion parameter of $\phi = 2$ (when $\sigma_{J_1} = 0.5$ and $\phi = 2$ the beta distribution is uniform between 0 and 1). Using different values of $\phi$ does not qualitatively change the existence of distinct extinction dynamics between populations with $\sigma_{A_R} < 0.5$ and those with $\sigma_{A_R} > 0.5$. However, increasing yearly variability in $\sigma_{J_1}$ increases extinction risk for all populations. For example, if we set $\phi = 0.001$, such that in a given year essentially either all year-1 juveniles survive or all of them die, populations with $\sigma_{A_R} > 0.5$ need to have $\sigma_{J_1}$ greater than 0.2 to have a 50-year extinction probability of less 50%. Because we do not pit tag juveniles, we do not have CMR estimates for $\sigma_{J_1}$ or $\phi$. However, based on qualitative and semi-quantitative field observations over 25 years, a value of $\sigma_{J_1} = 0.25$ in the presence of Bd is probably a reasonable estimate for many populations. Thus, we expect our model predictions to be conservative with regards to population recovery.

Finally, we assessed whether our stochastic model could reproduce observed trajectories of population recovery. We focused on population 70550 because this was our longest CMR time series for a translocated population and because this population shows evidence of substantial post-translocation increases in adult abundance associated with population establishment and recovery. We simulated our model for 16 years, repeating the simulation 50,000 times. For each run and each year, we drew $\sigma_{J_1}$ from a uniform distribution between 0 and 1 (or equivalently a beta distribution with mean 0.5 and $\phi = 2$). Using Approximate Bayesian Computing and rejection sampling (12), we identified the top 2% of trajectories (i.e., 1000 trajectories) that minimized the sum of squared errors between the observed and predicted data. The yearly $\sigma_{J_1}$ values associated with these "best" trajectories represented an approximate posterior distribution (13). Using these best fit trajectories, we assessed whether our model could qualitatively describe the patterns of recovery in the observed data for population 70550.

### 3. Frog evolution in response to Bd

**A. Study design.** To gain insights into the role of evolution in the development of resistance by MYL frogs, we compared frog exomes sampled in naive versus recovering populations. Comparing populations with different infection histories allowed larger sample sizes and replication across the landscape. The alternative approach of comparing samples from the same populations before and after Bd exposure isn't feasible in this system because Bd arrived in most MYL frog populations decades ago and population persistence/recovery is rare and unpredictable. As a result, samples from recovering populations collected before and after Bd exposure are not available and are unlikely to be available in the future.

Our study design, in which we compared frog genomes in naive and recovering populations, required sampling populations across the range of MYL frogs in the southern Sierra Nevada. This is due to the fact that very few MYL frog populations remain in the naive state, and those that do are scattered across a wide latitudinal range, from Yosemite National Park in the north to southern Kings Canyon National Park in south. To minimize potential confounding effects caused by known variation in frog genotypes across latitude (14), we selected sampling sites such that both population types were represented across similar latitudinal ranges (Fig. 5).

**B. GO analysis.** In **Results-Frog evolution in response to Bd**, we describe the stringent set of outlier variants (identified using a Bonferroni-corrected p-value of 0.01). A liberal set of outlier variants, identified using a Bonferroni-corrected p-value of 0.05, included 38 outliers (35 SNPs and 3 INDELS) from 30 distinct genes across 16 contigs. We used this liberal set to determine if any GO biological functions, molecular functions, or cellular processes were overrepresented. To do this, we retrieved the BLAST hits and mapped GO terms for each gene in our targeted transcriptome. We then conducted a statistical overrepresentation test (Fisher's exact test) using Blast2GO (15) to compare the 30 unique outlier genes to the complete set of genes in our target transcriptome. We repeated this process for the set of 35 genes located in the 9 shared regions of the $F_{ST}$ and $\pi_{diff}$ splined windows.

**C. Genetic diversity.** To characterize general patterns of genetic diversity between naive and recovering populations, we conducted three analyses. First, we calculated heterozygosity for each sampled frog using VCFtools (16). Second, to characterize genome-wide patterns of diversity, we used VCFtools to calculate nucleotide diversity ($\pi$) in 100kb sliding windows along the genome for each population. Third, we calculated average $\pi$ per population within each of the 9 outlier windows identified in the splined window analysis.

Average individual-level heterozygosity and genome-wide population-level $\pi$ were similar between the naive and recovering groups (Figure S8; Figure S9). Within each of the 9 outlier windows, average $\pi$ shows considerable variation between populations (Figure S10) and no obvious patterns between the naive and recovering groups.

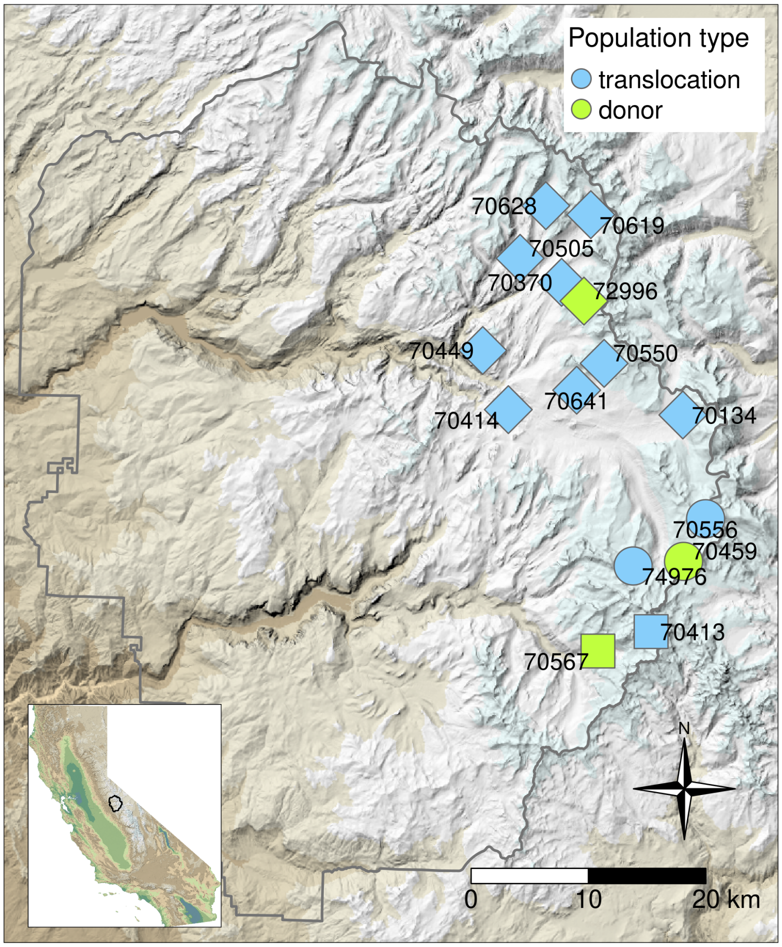


**Fig. S1.** Map showing the locations of translocated and donor MYL frog populations in Yosemite National Park (park boundary indicated by gray polygon). Symbol shapes indicate the donor population used for each translocation site, and 5-digit numbers identify each donor and translocation site. To obscure the exact locations of populations, random noise was added to all point coordinates. Inset map shows the location of Yosemite within California. In both maps, elevation is indicated by the colored hillshade layer (dark green = lowest elevation, white = highest elevation).

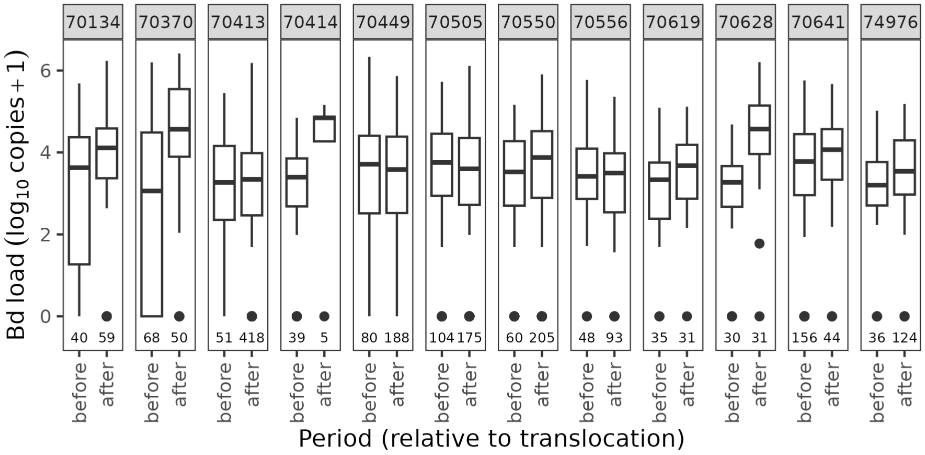


**Fig. S2.** For frogs translocated to each of the 12 recipient sites, Bd loads for the period immediately prior to translocation versus during the 1-year period after translocation. Bd loads are expressed as the number of ITS1 copies per skin swab, as estimated by qPCR of the Bd ITS1 region. Box plots show medians, first and third quartiles, largest and smallest values within 1.5x interquartile range, and values outside the 1.5x interquartile range. Loads indicative of severe disease are > 5.8 ITS copies (on a $log_{10}$ scale). Samples sizes are provided immediately above the x-axis.

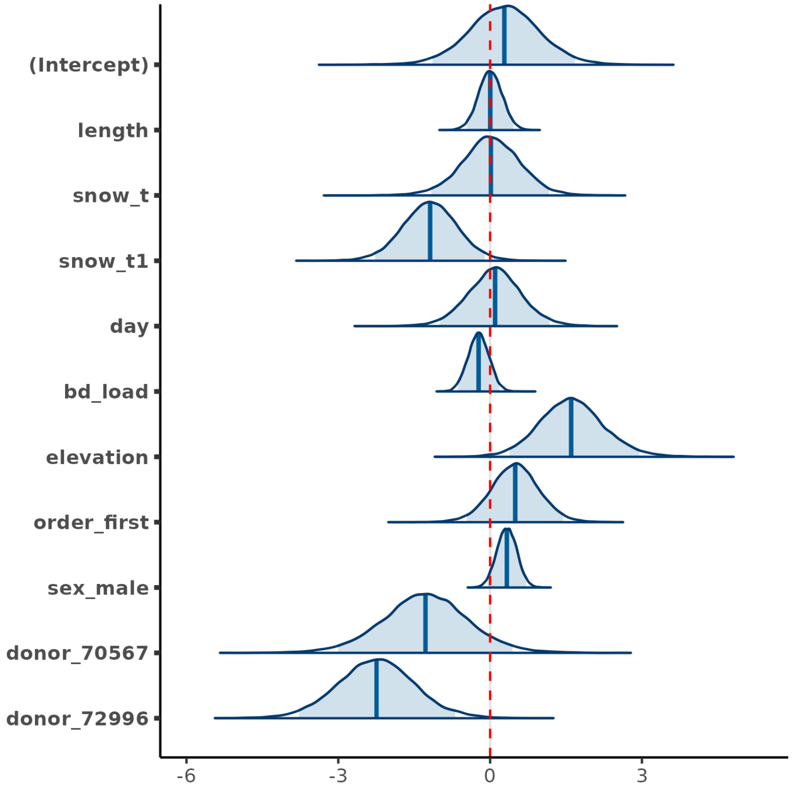


**Fig. S3.** Results from the among-site meta-analysis, showing that Bd load is not an important predictor of post-translocation frog survival. Depicted distributions are the estimated posterior density curves and shaded 95% uncertainty intervals for the intercept and all predictor variables from the best model. In the Bayesian framework in which the model was developed, variables are considered important predictors if the associated uncertainty interval does not overlap zero (indicated by the dashed red line). Predictor variables shown on the y-axis are defined as follows: length = frog size, snow_t = winter severity in the year of translocation (measured on April 1), snow_t1 = winter severity in the year following translocation (measured on April 1), day = day of year on which a translocation was conducted, bd_load = Bd load, elevation = site elevation, order_first = within-site translocation order, sex_male = frog sex, donor_70567 and donor_72996 = donor population.

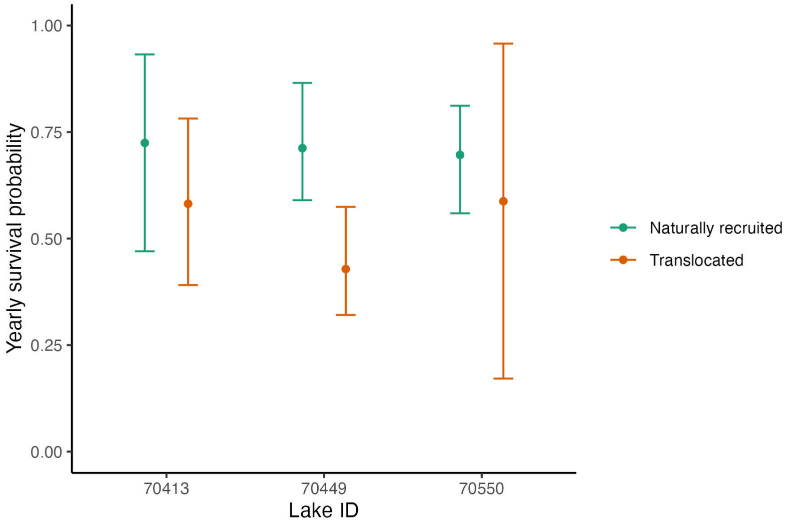


**Fig. S4.** Comparison of average yearly adult survival probabilities for adults translocated to each of 3 sites versus adults that were naturally recruited at each site (as a result of reproduction by translocated frogs). In contrast to Fig. 2, these are not survival probabilities from the first year following translocation, but instead represent averaged survival probabilities across multiple years and cohorts. Points are median estimates and error bars give the 95% uncertainty intervals around the estimates, accounting for yearly variation in survival probabilities. All estimates were derived using the mrmr package, and the methods for calculation are described in **Supporting Information - Population viability modeling - Estimating model parameters**.

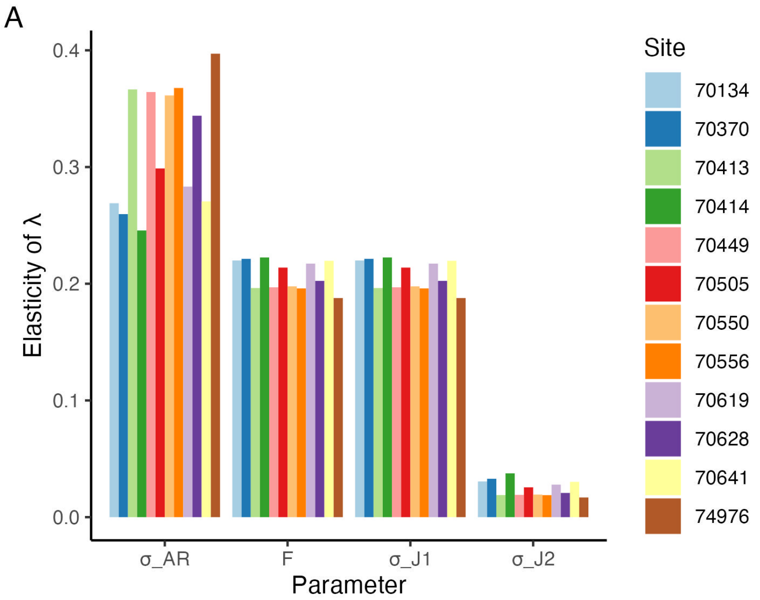


**Fig. S5.** Sensitivity analysis of the stage-structured MYL frog model. Elasticity of $\lambda$ with changes in four parameters: $\sigma_{A_R}$ (yearly survival probability of naturally recruited adults), *F (*number of eggs produced by a female frog in a year that successfully hatch), $\sigma_{J_1}$ (yearly probability of survival of year-1 juveniles that also affects recruitment), and $\sigma_{J_2}$ (yearly probability of survival and recruitment of year-2 juveniles). Elasticity is calculated at the default parameter values for each population and $\sigma_{J_1}$ = 0.09.

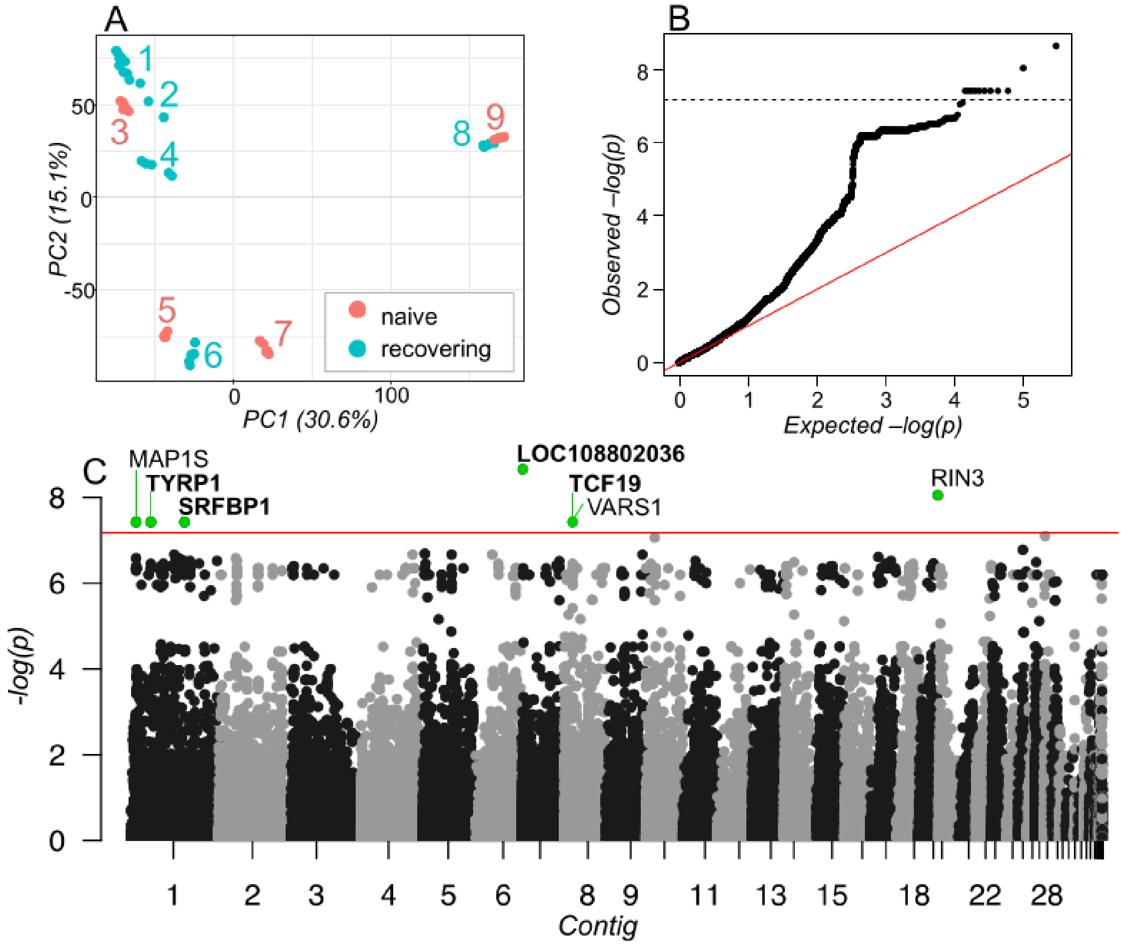


**Fig. S6.** Results from the analysis of individual variants, showing putative signatures of selection in recovering MYL frog populations. (A) PCA calculated from binary SNPs showing the genomic relationship of samples. Numeric labels and colors match those in Fig. 5. Populations 1-7 are *R. sierrae* and populations 8 and 9 are *R. muscosa.* (B) qqplot showing observed and expected p-values for 148,307 SNPs and INDELS. Dashed line shows p-value that identifies outliers. (C) Manhattan plot showing p-value for each SNP. SNPs are sorted by genomic position and contigs are sorted by size. Red line shows p-value that identifies outliers. Outlier SNPs above this threshold are highlighted and labeled. Bold labels indicate the presence of at least one non-synonymous SNP in that gene.

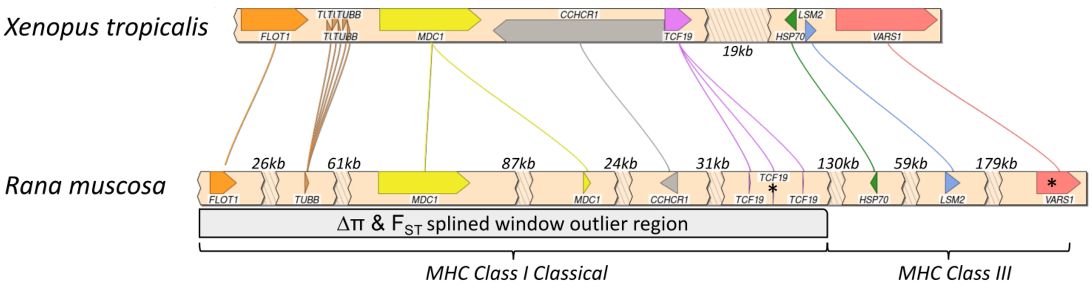


**Fig. S7.** Synteny plot showing conserved gene order in *Xenopus tropicalis* and *Rana muscosa* for the outlier region containing MHC Class I Classical and MHC Class III gene regions. The plot was created with SimpleSynteny (17) using *Xenopus tropicalis* Chromosome 8 (NC_030684.2, genbank accession GCA_000004195.4) and *Rana muscosa* Contig19. Asterices indicate the location of SNP outliers in the TCF19 and VARS1 genes. Gap sizes for each contig representation are labeled.

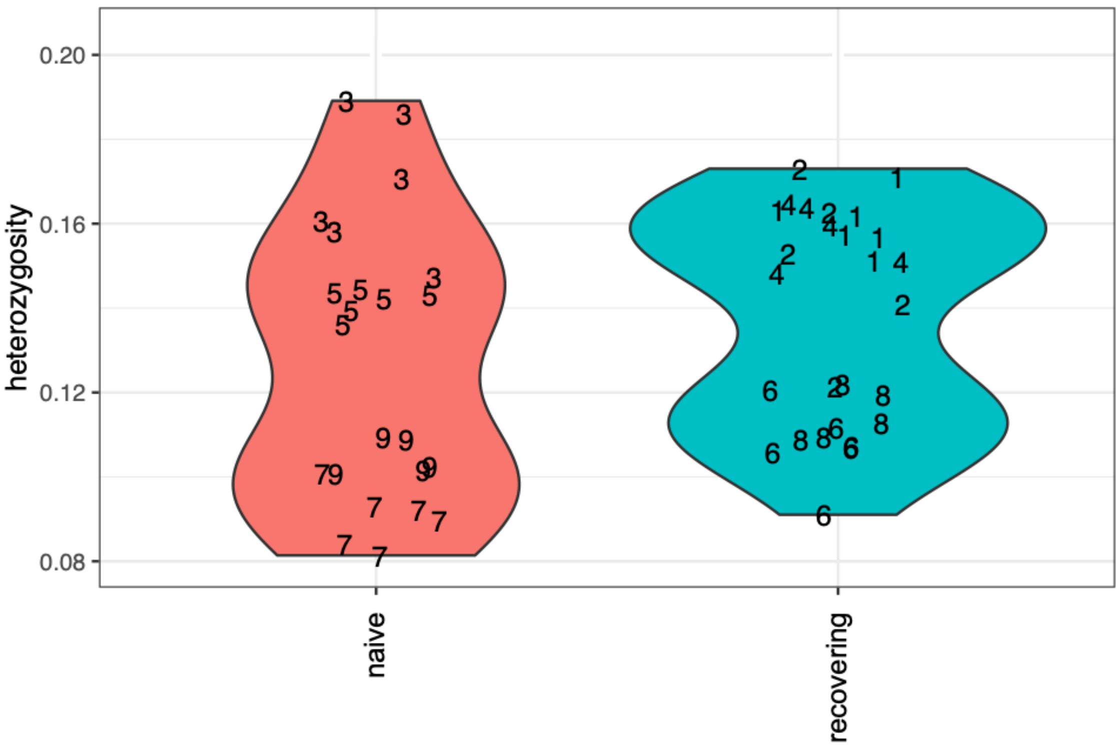


**Fig. S8.** Violin plots showing individual heterozygosity for the Bd-naive and recovering populations included in the frog evolution study. Individual data points are represented by their corresponding site number (from Fig. 5).

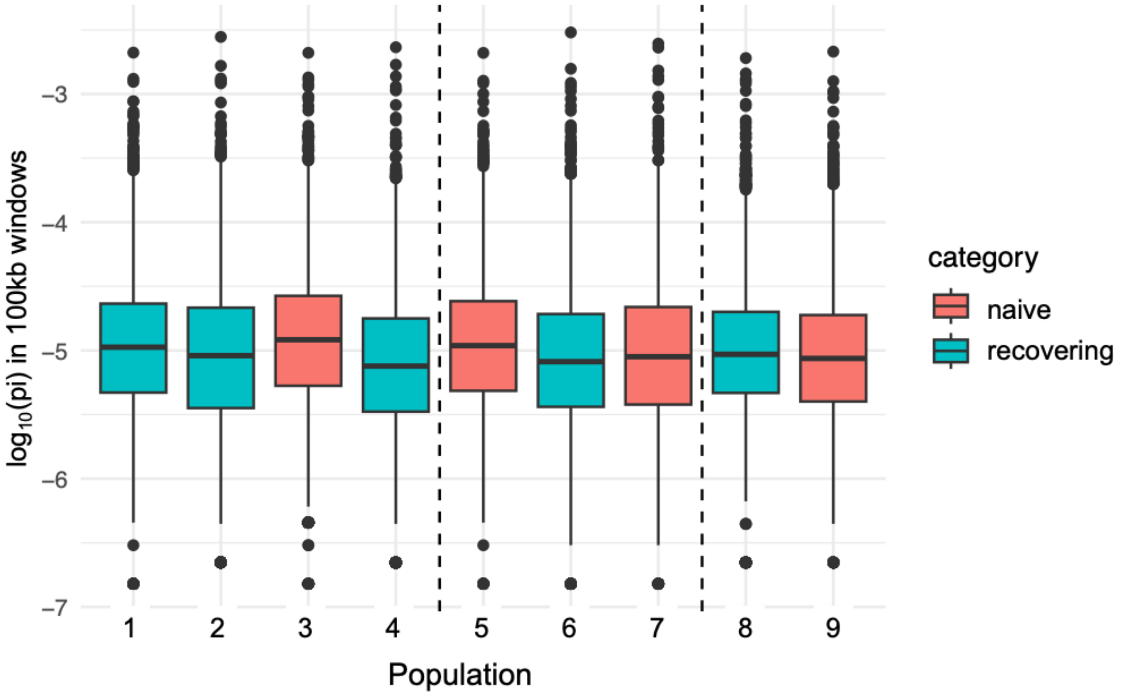


**Fig. S9.** For each study population in the frog evolution study (from Fig. 5), boxplots showing nucleotide diversity ($\pi$) within 100kb sliding windows along the genome. The y-axis has been $\log_{10}$-transformed for display purposes. Sites are color-coded as "Bd-naive" or "recovering", and vertical dashed lines show the genetic breaks between frog populations as described in Figure S6 A).

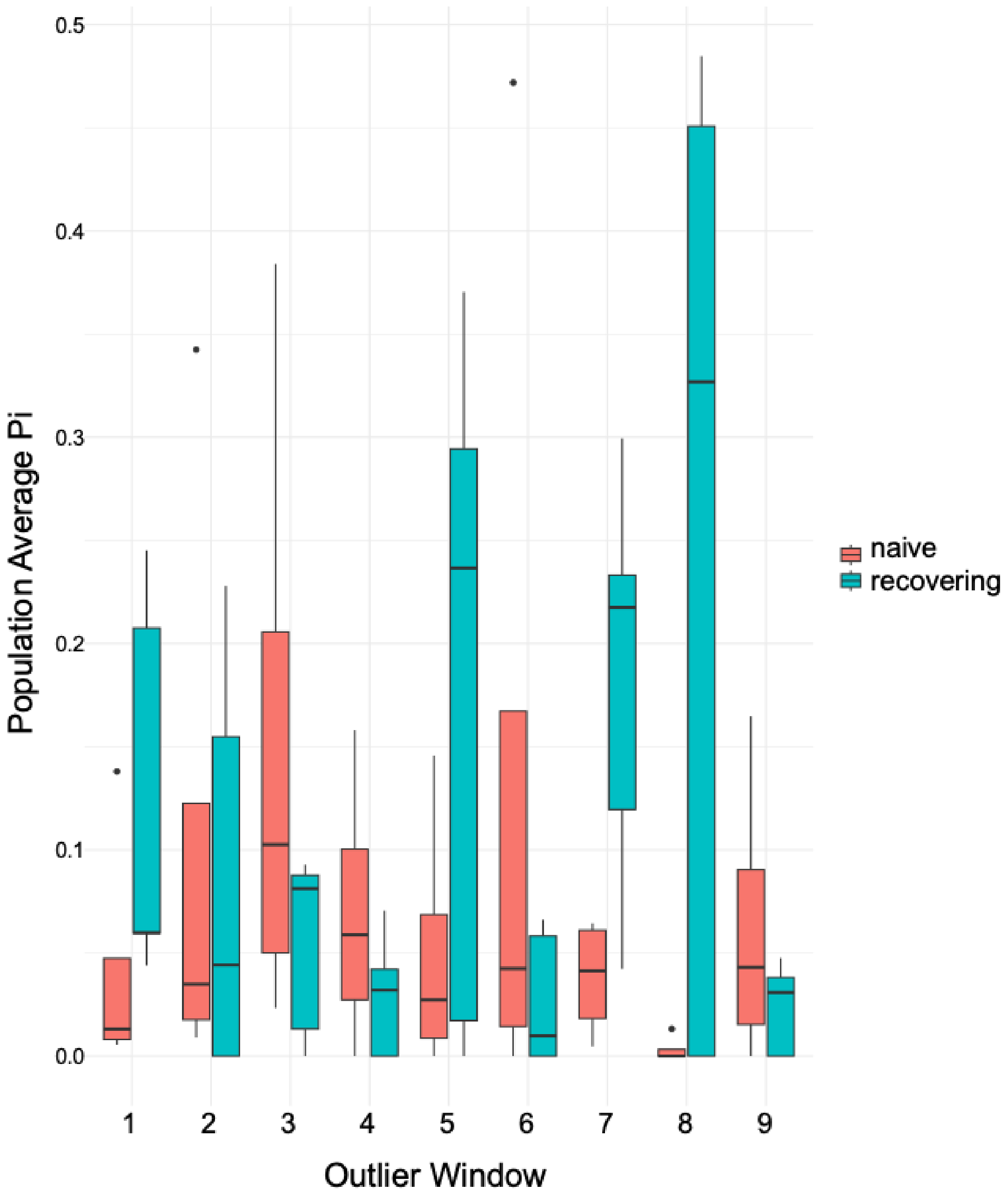


**Fig. S10.** For each of the 9 outlier windows identified by the overlapping splined windows for FST and $\pi$, boxplots showing average $\pi$ for naive and recovering populations. The corresponding gene annotations for each window are as follows: (1) C6, C7, (2) DDX10, ZBTB24, (3) SULTR-like, TRAF3IP2, VGLL3, EXOC1, (4) FLOT1, TUBB, MDC1, CCHCR1, TCF19, HSP70, LSM2, VARS1, (5) GCC2, CFAP251, PEG10, (6) ERO1A, GVINP1, (7) PPP1R12A, TSPAN4, PAWR, MFRP, MAX, PPP6R3, (8) C6H5ORF22, PKS6, BSPRY, MPV17, and (9) CAD, ATRAID, GPN1.

### C.1. Tables.

**Table S1. Description and values of parameters used in the model. All survival probabilities are in the presence of the fungal pathogen Bd.**

| Parameter | Value | Source |
|---|---|---|
| $\sigma_{L_1}$, Yearly survival probability of year-1 tadpoles | 0.7 | Estimated from field data, observations, natural history knowledge |
| $\sigma_{L_2}$, Yearly survival probability of year-2 tadpoles | 0.7 | Estimated from field data, observations, natural history knowledge |
| $\sigma_{L_3}$, Yearly survival probability of year-3 tadpoles | 0.7 | Estimated from field data, observations, natural history knowledge |
| $\sigma_{J_1}$, Yearly survival probability of year-1 juveniles | Varies yearly | Varies. Bounds estimated from field data, observations, natural history knowledge |
| $\sigma_{J_2}$, Yearly survival probability of year-2 juveniles | 0.5 | Estimated from field data, observations, natural history knowledge |
| $\sigma_{A_R}$, Yearly survival probability of naturally recruited adults | Varies by population | Estimated from CMR studies |
| $\sigma_{A_T}$, Yearly survival probability of translocated adults | Varies by population | Estimated from CMR studies |
| $p_{L_1}$, Probability of a year-1 tadpoles remaining as a tadpoles | 1 | Estimated from field data, observations, natural history knowledge |
| $p_{L_2}$, Probability of a year-2 tadpoles remaining as a tadpoles | 0.25 | Estimated from field data, observations, natural history knowledge |
| $p_{J_1}$, Probability of a year-1 juvenile remaining as a juvenile | 0.25 | Estimated from field data, observations, natural history knowledge |
| $p_F$, Probability of a adult female reproducing in a year | 0.5 | Could be as high at 1, based on field observations |
| $F$, Number of surviving eggs produced by an adult female | 100 | From observations of captive frogs |

**SI Dataset S1 (gemma_outliers_all.csv)**

Set of liberal SNP and INDEL outlier variants (Bonferroni corrected p-value $< 0.05$), identified via GEMMA.

**SI Dataset S2 (gemma_outliers_strict_freq.csv)**

Set of strict SNP and INDEL outlier variants (Bonferroni corrected p-value $< 0.01$), identified via GEMMA. Additional information includes variant location within the gene (predicted_gene_loc) and whether the variant is synonymous or nonsynonymous (predicted_effect_AA).

**SI Dataset S3 (spline_window_shared_outliers.csv)**

Description of overlapping $F_{ST}$ and $\pi_{diff}$ outlier windows, as identified in the splined window analysis.

**SI Dataset S4 (spline_window_gene_details.csv)**

Annotation information for all genes within each of the overlapping outlier windows in dataset S3

### References


1. D Boyle, D Boyle, V Olsen, J Morgan, A Hyatt, Rapid quantitative detection of chytridiomycosis (*Batrachochytrium dendrobatidis*) in amphibian samples using real-time Taqman PCR assay. *Dis. Aquatic Org.* **60**, 141–148 (2004).
2. K Kriger, J Hero, K Ashton, Cost efficiency in the detection of chytridiomycosis using PCR assay. *Dis. Aquatic Org.* **71**, 149–154 (2006).
3. AV Longo, D Rodriguez, D da Silva Leite, L Toledo, C Mendoza Almeralla, PA Burrowes, KR Zamudio, ITS1 copy number varies among *Batrachochytrium dendrobatidis* strains: implications for qPCR estimates of infection intensity from field-collected amphibian skin swabs. *PLoS ONE* **8**, e59499 (2013).
4. VT Vredenburg, RA Knapp, TS Tunstall, CJ Briggs, Dynamics of an emerging disease drive large-scale amphibian population extinctions. *Proc. Natl. Acad. Sci. USA* **107**, 9689–9694 (2010).
5. MB Joseph, RA Knapp, Disease and climate effects on individuals drive post-reintroduction population dynamics of an endangered amphibian. *Ecosphere* **9**, e02499 (2018).
6. JA Royle, RM Dorazio, Parameter-expanded data augmentation for Bayesian analysis of capture–recapture models. *J. Ornithol.* **152**, 521–537 (2012).
7. W Zucchini, IL MacDonald, *Hidden Markov models for time series: an introduction using R.* (Chapman and Hall/CRC), (2009).
8. MB Joseph, mrmr: mark recapture miscellany in R (2019) R package version 0.1.1.
9. B Goodrich, J Gabry, I Ali, S Brilleman, rstanarm: Bayesian applied regression modeling via Stan. (2022) R package version 2.21.3.
10. D Lewandowski, D Kurowicka, H Joe, Generating random correlation matrices based on vines and extended onion method. *J. Multivar. Analysis* **100**, 1989–2001 (2009).
11. A Vehtari, A Gelman, J Gabry, Practical Bayesian model evaluation using leave-one-out cross-validation and WAIC. *Stat. Comput.* **27**, 1413–1432 (2016).
12. M Kosmala, P Miller, S Ferreira, P Funston, D Keet, C Packer, Estimating wildlife disease dynamics in complex systems using an Approximate Bayesian Computation framework. *Ecol. Appl.* **26**, 295–308 (2016).
13. MA Beaumont, Approximate Bayesian Computation in evolution and ecology. *Annu. Rev. Ecol. Evol. Syst.* **41**, 379–406 (2010).
14. AQ Byrne, AP Rothstein, LL Smith, H Kania, RA Knapp, DM Boiano, CJ Briggs, AR Backlin, RN Fisher, EB Rosenblum, Revisiting conservation units for the endangered mountain yellow-legged frog species complex (*Rana muscosa*, *Rana sierrae*) using multiple genomic methods. *Conserv. Genet.* (2023).
15. A Conesa, S Götz, JM García-Gómez, J Terol, M Talón, M Robles, Blast2GO: a universal tool for annotation, visualization and analysis in functional genomics research. *Bioinformatics* **21**, 3674–3676 (2005).
16. P Danecek, A Auton, G Abecasis, CA Albers, E Banks, MA DePristo, RE Handsaker, G Lunter, GT Marth, ST Sherry, G McVean, R Durbin, The variant call format and VCFtools. *Bioinformatics* **27**, 2156–2158 (2011).
17. D Veltri, MM Wight, JA Crouch, Simplesynteny: a web-based tool for visualization of microsynteny across multiple species. *Nucleic Acids Res.* **44**, W41–W45 (2016).
